# The biophysical basis of bacterial colony growth

**DOI:** 10.1101/2023.11.17.567592

**Authors:** Aawaz R. Pokhrel, Gabi Steinbach, Adam Krueger, Thomas C. Day, Julianne Tijani, Siu Lung Ng, Brian K. Hammer, Peter J. Yunker

## Abstract

Bacteria often attach to surfaces and grow densely-packed communities called biofilms. As biofilms grow, they expand across the surface, increasing their surface area and access to nutrients. Thus, the overall growth rate of a biofilm is directly dependent on its “range expansion” rate. One factor that limits the range expansion rate is vertical growth; at the biofilm edge there is a direct trade-off between horizontal and vertical growth—the more a biofilm grows up, the less it can grow out. Thus, the balance of horizontal and vertical growth impacts the range expansion rate and, crucially, the overall biofilm growth rate. However, the biophysical connection between horizontal and vertical growth remains poorly understood, due in large part to difficulty in resolving biofilm shape with sufficient spatial and temporal resolution from small length scales to macroscopic sizes. Here, we experimentally show that the horizontal expansion rate of bacterial colonies is controlled by the contact angle at the biofilm edge. Using white light interferometry, we measure the three-dimensional surface morphology of growing colonies, and find that small colonies are surprisingly well-described as spherical caps. At later times, nutrient diffusion and uptake prevent the tall colony center from growing exponentially. However, the colony edge always has a region short enough to grow exponentially; the size and shape of this region, characterized by its contact angle, along with cellular doubling time, determines the range expansion rate. We found that the geometry of the exponentially growing biofilm edge is well-described as a spherical-cap-napkin-ring, i.e., a spherical cap with a cylindrical hole in its center (where the biofilm is too tall to grow exponentially). We derive an exact expression for the spherical-cap-napkin-ring-based range expansion rate; further, to first order, the expansion rate only depends on the colony contact angle, the thickness of the exponentially growing region, and the cellular doubling time. We experimentally validate both of these expressions. In line with our theoretical predictions, we find that biofilms with long cellular doubling times and small contact angles do in fact grow faster than biofilms with short cellular doubling times and large contact angles. Accordingly, sensitivity analysis shows that biofilm growth rates are more sensitive to their contact angles than to their cellular growth rates. Thus, to understand the fitness of a growing biofilm, one must account for its shape, not just its cellular doubling time.

## 1. Introduction

Biofilms - surface attached communities of microbes embedded in extracellular polymeric substances - represent a major mode of microbial life (1–4). During the biofilm life cycle, cells attach to a surface, secrete extracellular polymeric substances, and reproduce, forming a biofilm that expands as it grows (5). Biofilm growth is limited by the ability of densely packed cells to access nutrients; expansion that increases a biofilm’s interface with nutrient-providing media thus enables more growth. Higher range expansion rates, i.e., the horizontal growth rate of a biofilm, therefore correlate with higher fitness. However, while biofilms grow they can expand horizontally and vertically (6–9), and at the edge of a colony there is a direct trade-off between horizontal and vertical growth (i.e., the more a biofilm grows vertically, the less it can grow horizontally). And yet, little is known about the impact of the biophysical balance between horizontal and vertical growth, despite the important role this trade-off plays in determining colony growth rate.

When biofilms are small, they typically grow exponentially (10). Once biofilms are tall enough, their growth is limited by nutrient diffusion and uptake and thus is sub-exponential (11–13). However, there is always a region at the biofilm edge that is thin enough to overcome the diffusion limitation and grow exponentially (7, 14–16). The range expansion rate is thus strongly dependent on the number and arrangement of cells in this region, which, in turn, depends on biofilm geometry. The biofilm geometry has also been demonstrated to significantly impact cell organization and packing which has been shown to impact the fitness of colony (10, 17–21). Determining the exact geometry of a biofilm and the edge, and how it changes over time, however, is experimentally challenging. Doing so requires high spatial and temporal resolution measurements of the three-dimensional biofilm surface topography across many length and time scales, which is experimentally difficult. Thus, we lack an empirical understanding of the shape of the edge, its time evolution, and how it relates to the range expansion rate.

Here, we examine three-dimensional biofilm geometries and how they impact biofilm growth by focusing on the shape and dynamics of the expanding colony edge. We find that expanding colonies rapidly approach consistent, steady-state contact angles upon growth. When small, biofilms are well-described as spherical caps. As biofilms grow larger, nutrient diffusion and uptake limit growth in the center of the biofilm, leading to deviations from spherical cap shapes; further, buckling and wrinkling may occur in the biofilm center (22). However, the contact angle at the edge remains constant. We show that the edge of the biofilm is well described as a spherical-cap-napkin-ring (SCNR), i.e., a spherical cap with a cylinder removed from its center. We derive expressions for the volume of SCNR and for the range expansion rate based on the doubling time of individual cells and the SCNR geometry. We measure growth of many strains, and some on different agar percentages, and show that differences in range expansion rate depend more on biofilm contact angle than on cellular doubling times.

## 2. Results

### A. Range expansion rates correlate more with colony geometry than with doubling time

As a first step, we sought to assess how much the range expansion rate depends on cellular reproduction and biofilm geometry. We measured three different properties: the lateral range expansion rate, Δ*a/*Δ*t*, where *a* is the colony radius and *t* is time, the cellular doubling time, *τ*, and the “contact angle,” the angle that the edge of the biofilm makes with the agar surface, *θ*, which characterizes the balance between vertical and horizontal growth (23). Experiments were done on five bacterial strains across three different species (*Pseudomonas aeruginosa, Aeromonas veronii, Vibrio cholerae wild type (WT), V. cholerae EPS-*, and *V. cholerae EPS+*; see Fig 1 and the Methods Section for more details on the strains used) on 1.5% LB agar pads . As we will primarily focus on *V. cholerae WT* in this paper, we also performed experiments with it on 3% and 5% agar pads. A total of seven independent strain-agar-concentration combinations were tested.

**Fig. 1.**
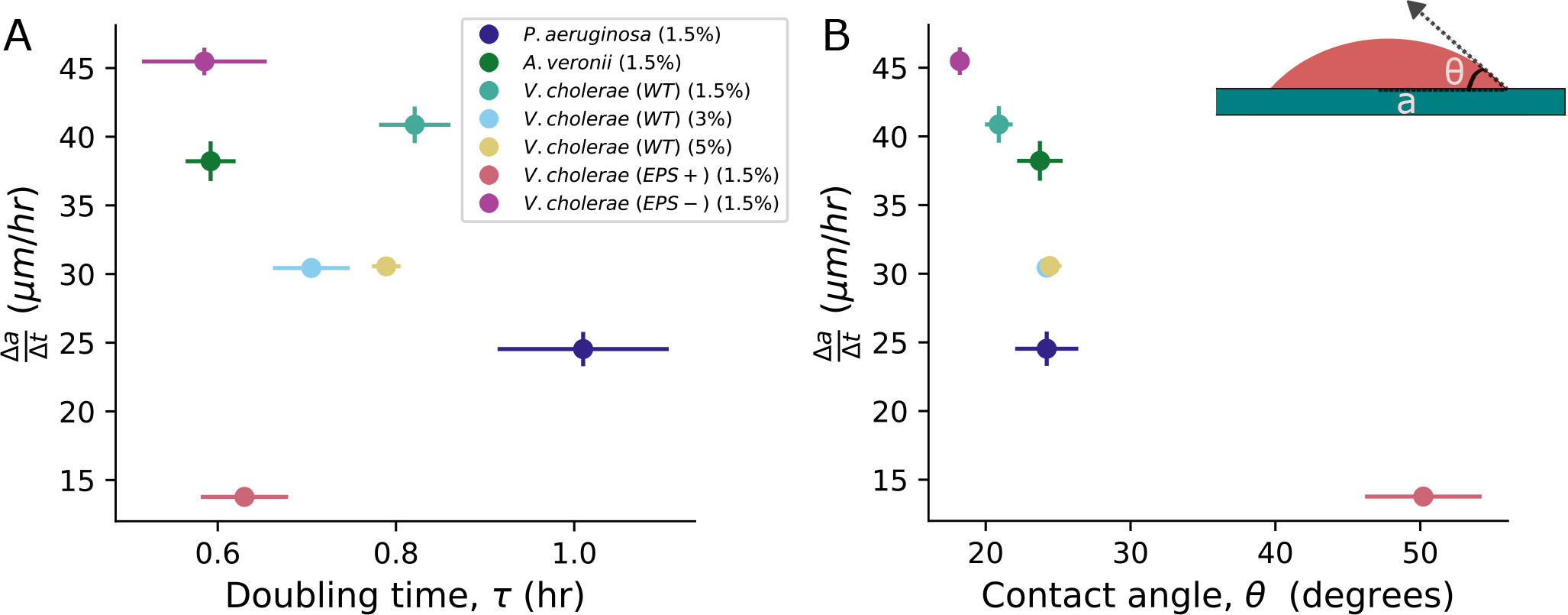
Dependence of range expansion rate (Δ*a/*Δ*t*) on doubling time (*τ* ) and contact angle (*θ*). **(A)** We grew monoclonal colonies at 23° C and, after 24 hours, measured the radius at four distinct time points to determine Δ*a/*Δ*t* for each colony. We then plotted the range expansion rate against the mean *τ* of the bacteria, directly measured from confocal microscopy with single cell resolution. The resulting Pearson’s correlation coefficient (*p* = −0.23) suggests a relatively weak correlation between these two parameters. The error bars in *τ* represent the standard deviation measured across at least three distinct replicates of bacteria, while the error bars in Δ*a/*Δ*t* represent the error in the mean slope calculated from three replicates of the biofilm. Different color symbols represent different strains and conditions, as described in the inset; note, the number in parentheses refers to the agar percentage of the pad on which the bacteria were grown. **(B)** Conversely, when we plotted Δ*a/*Δ*t* against the mean *θ* of each colony, we observed a significant correlation (*p* = *−*0.85), suggesting that edge geometry plays a crucial role in range expansion. The error bars in *θ* represent the error in the mean angle calculated across three replicates.

Doubling times were measured by inoculating single cells on agar surfaces and observing their growth, from a single cell to ∼ 16 cells, via confocal microscopy (Nikon A1R). Range expansion rates and contact angles were measured by inoculating single cells on agar surfaces and measuring their topographies after ∼ 24 hours of growth with optical interferometry (Zygo ZeGage). With these measurements in hand, we characterized how Δ*a/*Δ*t* correlates with *τ* and *θ*. Surprisingly, the correlation between Δ*a/*Δ*t* and *τ* was relatively weak (Pearson’s correlation coefficient *ρ* = −0.23). Conversely, the correlation between Δ*a/*Δ*t* and *θ* was much stronger (*ρ* = −0.85). These observations suggest that colony geometry may play a crucial role in determining range expansion rates, and thus biofilm fitness.

### B. Nascent biofilm geometry is well described by a spherical cap

To investigate how colony geometry impacts range expansion rate, we next sought to characterize the full three-dimensional colony shapes. To do so, we inoculated single cells of *V. cholerae WT* on 1.5% LB agar pads. After growing colonies for ≈ 24 hours at 23° C, we used optical interferometry to measure the colony surface topography.

We observed that at these sizes, colonies look like spherical caps. Fig. 2A shows an image of a *V. cholerae WT* biofilm grown for ≈ 24 hrs on 1.5% agar projected onto a single plane (the X-Z plane). To test if this morphology is unique to *V. cholerae WT*, we performed similar experiments with five more strains, including gram-negative and gram-positive species and different cell shapes and sizes. We observed a spherical cap shape for all strains (Fig. 2B-F).

**Fig. 2.**
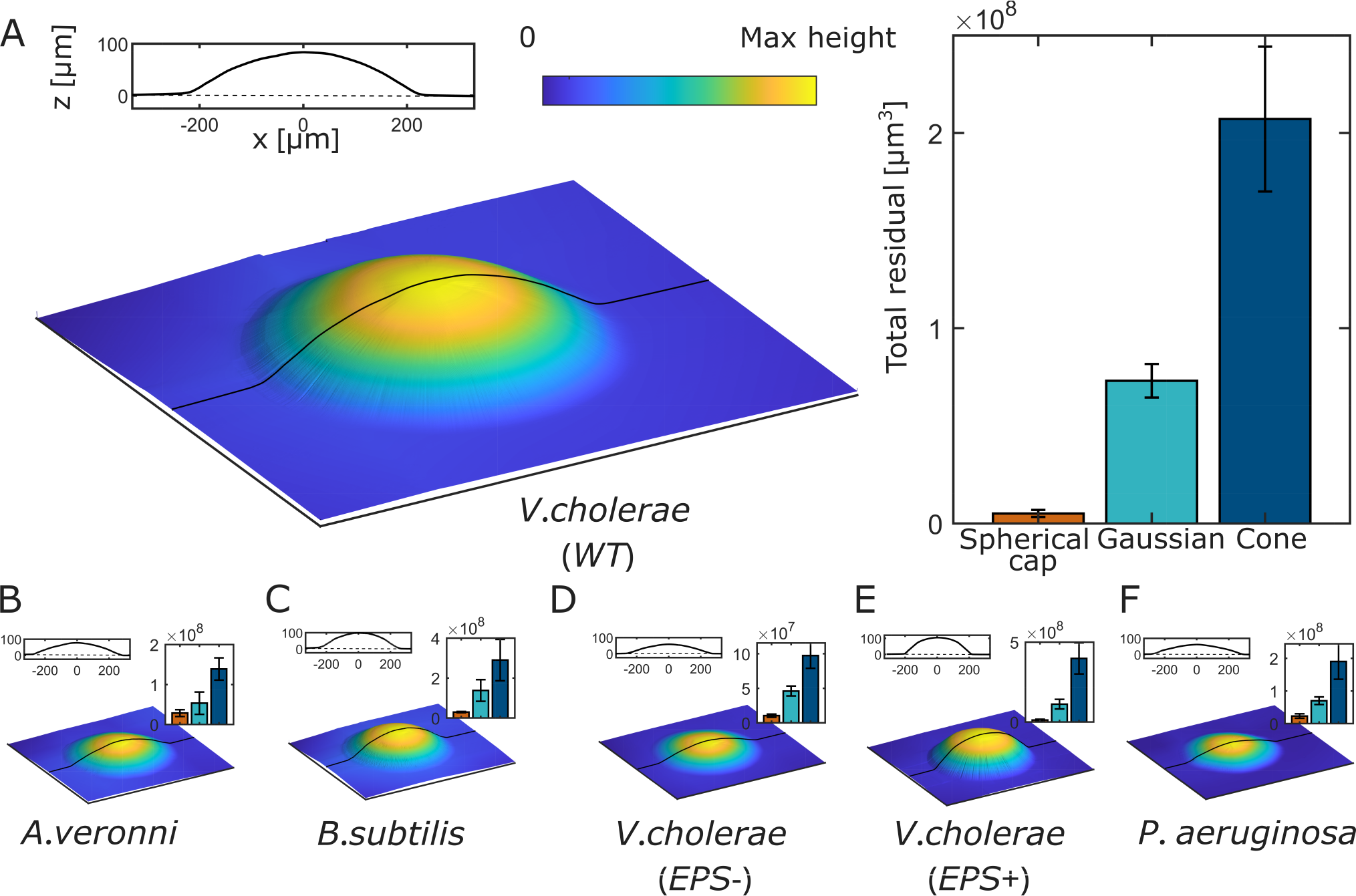
Topographical images of bacterial colonies. **(A)** The topography of a monoclonal colony of *V. cholerae WT* is measured via interferometry after *∼* 24 hours of growth on a 1.5% agar plate. The 2D line profile shows the topographic profile in the x-z plane. We compared this topography to a perfect spherical cap with experimentally measured height and radius, and did the same for Gaussian and cone shapes. Residuals from all three comparisons are shown in the bar plot, which demonstrates that the spherical cap shape yields by far the lowest residual. **(B-E)** We performed similar measurements on five additional strains: *Aeromonas veronii, Bacillus subtilis, V. cholerae EPS-, V. cholerae EPS+*, and *Pseudomonas aeruginosa*. Topographies, x-z plane topographic profiles, and residuals are shown for these strains as in (A). In all cases, the spherical cap produced the smallest residual, indicating the broad prevalence of the spherical cap shape.

We next sought to quantify how well these biofilms are described as spherical caps. First, we measured radius (*a*) and height (*h*) for each strain. These two measurements are sufficient to completely define a spherical cap. We then compared the empirically measured topographies to ideal spherical caps with the same *a* and *h*; for context, we compared to other possible shapes–specifically, the Gaussian and Cone. For all strains, we found an excellent agreement between measured topographies and their associated spherical caps, and much worse agreement with the Gaussian and Cone shapes, as quantified by the coefficient of determination, *R*^2^ (for all six strains, *R*^2^ *>* 0.95 for spherical caps, *R*^2^ *<* 0.9 for Gaussians, and *R*^2^ *<* 0.7 for cones; see SI Fig. S1). We also measured the total residuals from all three shapes (See SI for details). We found that the spherical cap always has by far the smallest residual, across all species and strains (Fig. 2B-E). Note, here we are not comparing the least-squares spherical cap fits, but simply with spherical caps with the same *a* and *h* as seen experimentally.

To further confirm that the spherical cap is indeed the best description of colony geometry, we calculated a least squares fit for spherical cap and Gaussian shapes for each topography (we omitted the cone due to its poor *R*^2^ and total residual). The free parameters obtained from least squares fit are in excellent agreement with the measured parameters for spherical cap shape (e.g., a plot of measured *R* vs. fit *R* has slope 0.91 and *R*^2^ = 0.99; see SI Fig. S2). In contrast, the measured and fit parameters do not match for Gaussian shapes (e.g., plots of measured vs. fit for *σ*_*X*_ and *σ*_*Y*_ have slopes -.003 and -0.003, and *R*^2^ = 0.4 and 0.5, respectively; see SI Fig. S3). As a final confirmation of the appropriateness of describing these colonies as spherical caps, we used the radius and height of each topography to calculate the volume of its corresponding spherical cap. These volumes match well with the experimentally measured volumes, with slope of 1.05 and *R*^2^ = 0.96 (See SI Fig. S5). Taken together, these data demonstrate that these nascent biofilms are well-described as spherical caps.

However, the spherical cap shape cannot persist forever. It is known that the vertical growth of a biofilm is limited by nutrient diffusion and uptake (7). The center of the biofilm eventually grows slower than the edges, modifying the process that produces a spherical cap. It is unclear how this vertical growth limitation affects geometry over time, and thus impacts the range expansion rate.

### C. Mature biofilms deviate from spherical cap shape

We next characterized how geometry changes over time. We grew *V. cholerae* WT colonies starting from a single cell on 1.5% LB agar. *≈* 17 hrs after inoculation, when the colony radius was ∼ 100 μm, we began measuring colony topography via interferometry. We imaged each colony every ≈ 15 min for the first 8 hours, then every ≈ 60 min for another 15 hours.

By visualizing the topographic profiles over time, we made two qualitative observations (Fig. 3A). First, while nascent biofilms are very well-described as spherical caps, mature biofilms increasingly deviate from spherical cap shapes. Second, the contact angle at the biofilm edge appears relatively constant across all times. Thus, we sought to quantify how the shape of a biofilm deviates from that of a spherical cap over time.

**Fig. 3.**
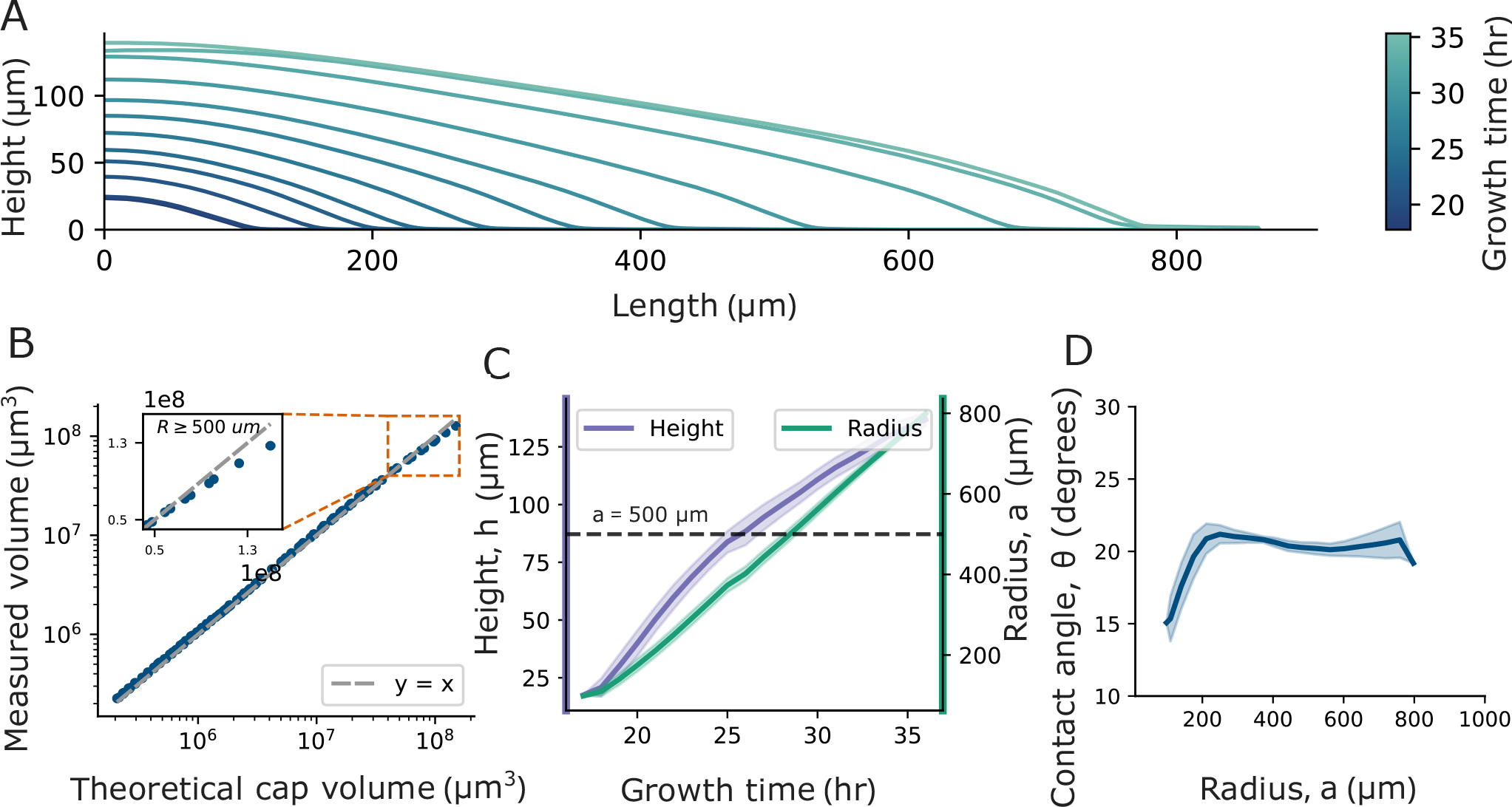
Change of colony geometry over time. **(A)** X-Z profiles of a *V. cholerae (WT)* colony measured at different times are shown; the time of each measurement is indicated by its color, as defined by the color bar. The colony initially exhibits a spherical cap shape, but over time its height saturates, and the colony gradually deviates from the spherical cap shape. **(B)** To quantitatively evaluate how the colony’s shape eventually deviates from the spherical cap shape, we compared the measured volume to the theoretical spherical cap volume, which we calculated using empirically measured values of the radius (*a*) and contact angle (*θ*). The measured volume and theoretical spherical cap volume start in good agreement. As the colony grows, the measured volume eventually falls short in comparison to the theoretical volume, demonstrating the deviation from the spherical cap shape. **(C)** We plotted the radius and height of the colony over time and found that while the radius grows linearly over time, the height of the colony increases at a slower rate after *a >* 500 μm. The solid line represents the LOESS-averaged values, and the shaded region represents the standard deviation from three replicates. The dashed line indicates *a* = 500 μm. **(D)** Contact angle, *θ*, is plotted against *a*, revealing that, after an initial increase when Δ*a/*Δ*t* is increasing, *θ* remains relatively constant.

We first measured volume (*V* ), *a*, and *θ* of colonies as functions of time. To determine when the biofilm deviates from a spherical cap shape, we compared *V* to the volume of a perfect spherical cap with the same *a* and *θ*, i.e., *V*_*cap*_(*a, θ*), which is given by

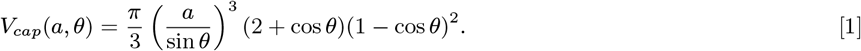

We found that *V* and *V*_*cap*_ are very similar until *a* ≈ 500 μm (Fig. 3B). At that point the percent difference between *V* and *V*_*cap*_ starts increasing and eventually exceeds 10%, and proceeds to increase further as *a* increases (see SI Fig. S6). Thus, while biofilms with *a <* 500 μm are well-described as spherical caps, those with *a >* 500 μm deviate from spherical cap geometries (similar trends were also observed for biofilms grown on different agar percentages, see SI Fig. S6).

To elucidate the cause of this deviation, we quantified the temporal evolution of all three parameters that characterize a spherical cap, i.e., *a, h*, and *θ* (a spherical cap is uniquely defined by any two of these three parameters). As expected, we observed that after an initial period of exponential growth, *a* increases linearly with time (Fig. 3 C and SI Fig. S8). In contrast, we found that *h* grows slower when *a >* 500 μm, as expected due to the effect of diffusion and uptake of nutrients (7) (Fig. 3C). However, unlike *a* and *h, θ* remains relatively constant after the initial period of exponential growth (Fig. 3D). This constant contact angle is not just limited to colonies grown on 1.5 % agar, but also in colonies grown on 3 and 5% agar (see SI Fig. S7 A). As the radius of the colony is growing at a constant rate, and the contact angle is constant, the saturation of *h* thus prevents the colony from maintaining a true spherical cap shape.

To confirm that the contact angle remains relatively constant despite the fact that colonies deviate from spherical cap shapes, we performed additional measurements of contact angle at late times. Even after 80 hours, when *a* = 1400 μm, *θ* = 40°; the mean contact angle fluctuates with a standard deviation of 1.4° from *a* = 300 μm to *a* = 1400 μm (which is the largest radius we measured; see SI Fig. S7B for details). So, even though biofilms cease to be spherical caps at late times, the geometry at the edge—where rapid growth occurs—does not change substantially even over long times. Further, we observe that biofilms that substantially wrinkle also exhibit clear, easily measurable contact angles at the biofilm edge (see SI Fig. S9; further, below we will show that the contact angle of biofilms that wrinkle has the same effect on their range expansion as it does for biofilms that do not wrinkle). Inspired by this observation, and with the ultimate goal of connecting accurate biofilm geometries to range expansion rates, we next sought to determine if the biofilm edge continues to be well-described as a spherical cap.

### D. The edge of a biofilm is described by Spherical Cap Napkin Ring Geometry

To determine if the biofilm edge continues to look like a spherical cap, we turned to the so-called Napkin Ring Problem. This is a well-known problem in geometry in which a cylindrical volume is removed from a sphere (the sphere and cylinder must share the same center) (24). Since our biofilms, when small, are well described as a spherical cap, and the edge retains a constant contact angle, we use a similar approach to a napkin ring problem to derive the volume of a “spherical cap napkin ring,” i.e., a spherical cap with a cylinder removed from its center (see Fig. 4 A and SI for details). We find that the volume of a spherical cap napkin ring is given by

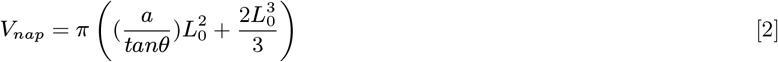

where *a, θ* and *L*_0_ are radius of the spherical cap, contact angle and edge height respectively.

**Fig. 4.**
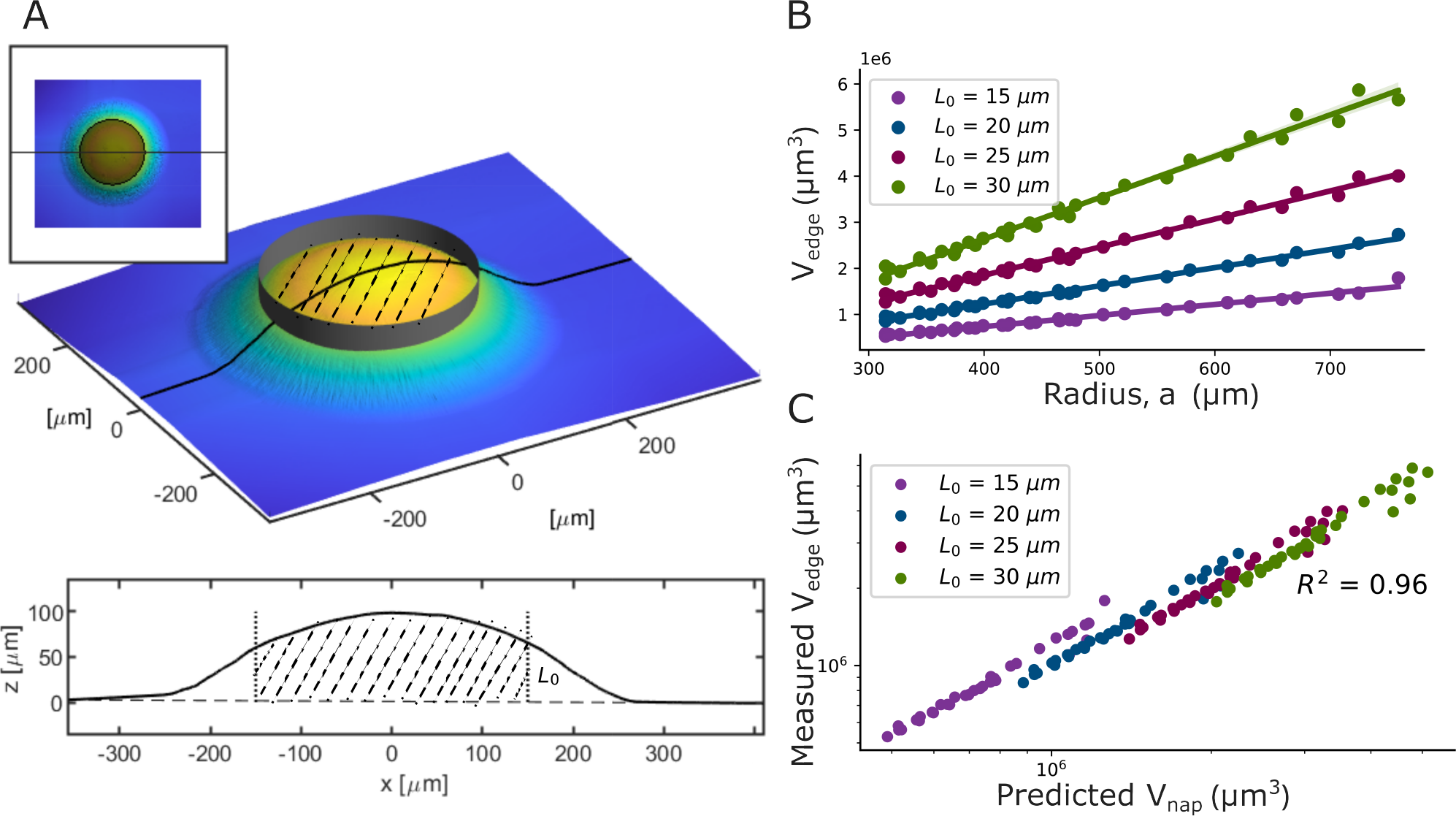
Definition and application of the Spherical-Cap-Napkin Ring geometry. **(A)** The sketches illustrate the process of removing the center from a spherical cap, thus defining a spherical-cap-napkin-ring, i.e., the material remaining at the cap’s edge. **(B)** We plotted the edge volume (*V*_edge_) against the colony radius for different edge heights (*L*_0_) for *V. cholerae (WT)* colony grown on 1.5% agar. Consistent with Equation 2, our derived expression for the volume of a spherical-cap-napkin-ring (See SI for derivation details), in the regime where the contact angle is constant, the edge volume increases linearly with the colony radius (*R*^2^ *>* 0.9 for all *L*_*e*_). The shaded region indicates the 95% confidence interval for the linear fit. **(C)** We plot the measured (*V*_edge_) against the predicted volume from Eq. 2, *V*_*nap*_. The high value of *R*^2^ = 0.96 demonstrates excellent agreement between the measured and predicted volumes.

We next experimentally check if real biofilm edges are accurately described as spherical cap napkin rings. First, we note that the volume of a spherical cap napkin ring depends on the spherical cap radius (*a*), contact angle (*θ*) and the edge “height” (*L*_0_). In experiments, we observe that, in the regime where contact angle is constant, the volume of the biofilm edge increases linearly with *a* for a wide range of value of *L*_0_ (Fig. 4B and SI Fig. S12A, *R*^2^ *>* 0.9 for all *L*_0_). Further, we find that the volume of the edge of a biofilm strongly agrees with predictions from the spherical cap napkin ring formula. In fact, in the regime where *θ* is constant (*a >* 300*μm*), measured volume of the edge versus predicted napking ring volume has an *R*^2^ = 0.96 with a slope of 1.0 (Fig. 4C and SI Fig. S12 B ). Thus, the geometry at the edge of a biofilm is, in fact, well-described as a spherical cap napkin ring.

### E. Consequences of the Spherical Cap Napkin Ring Geometry

We now use the SCNR geometry to predict the range expansion rate. We first sought a simpler approach to capture the impact of the geometry at the biofilm edge on the range expansion rate. We derived an expression for the change in radius as a function of time for a spherical cap napkin ring, building off the spherical cap napkin ring volume from Eqn.2; to leading order, we have:

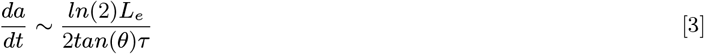

where *L*_*e*_ represents the maximum height at the edge of the biofilm where the volume doubles with the doubling time of *τ* (see SI for more details). In fact, we find that experimentally measured Δ*a/*Δ*t* exhibits a very strong correlation with ∼ 1*/*(*τtan*(*θ*)), with a correlation coefficient of 0.93 (Fig. 5A). Furthermore, the linear relationship in Eq 3 allows us to extract a value of *L*_*e*_. The best linear fit to these data, as shown in Figure 5A, gives *L*_*e*_ = 29*μm*.

**Fig. 5.**
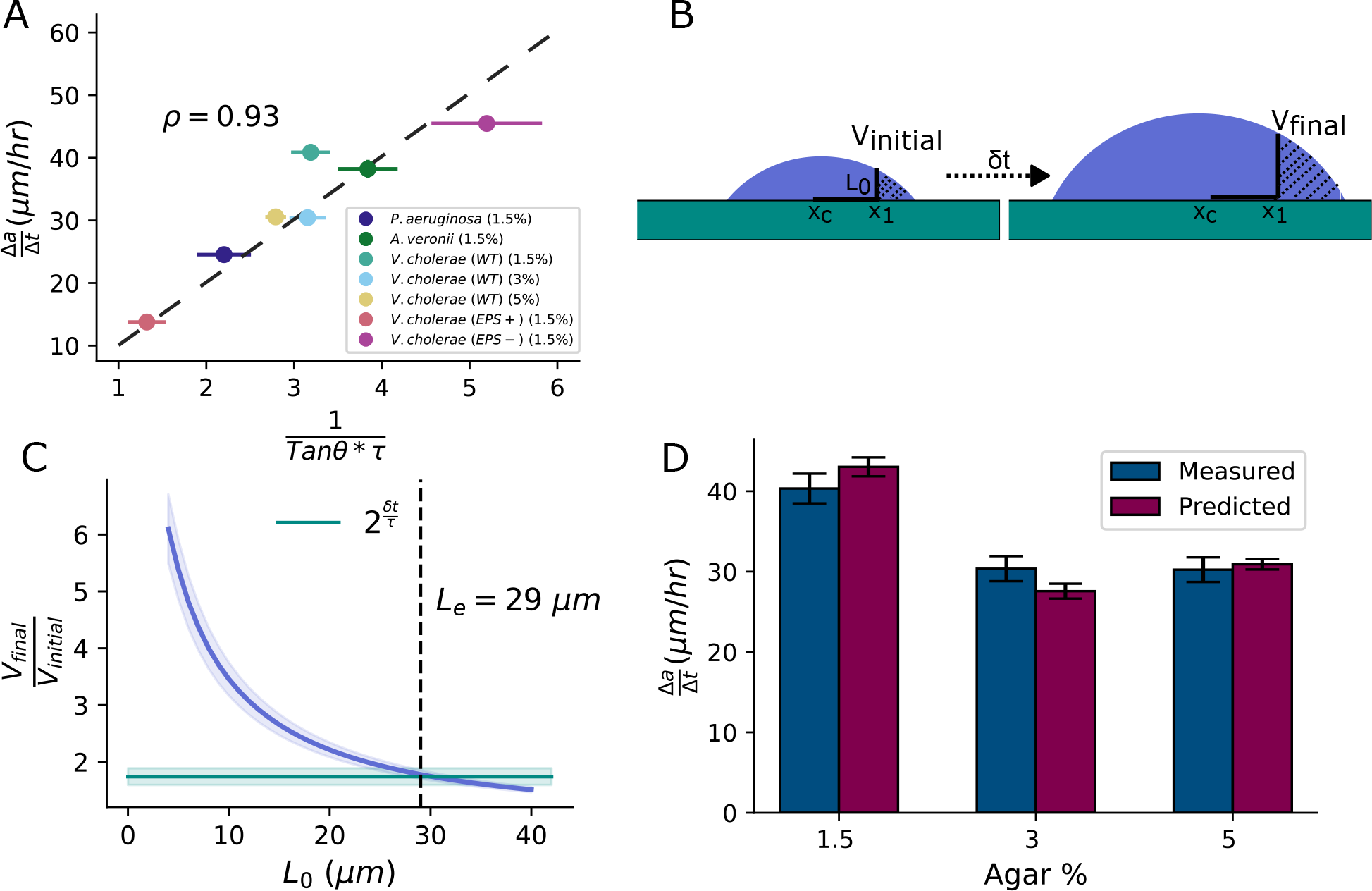
Experimental measurements agree with predictions from the spherical cap napkin ring model. **(A)** The range expansion rate (Δa/Δt) is plotted against ∼ 1*/*(*τ* tan(*θ*)). In line with the spherical-cap-napkin-ring predictions, we found a strong Pearson correlation of *ρ* = 0.93. Using Equation 3, measurement of the slope allows us to estimate that *L*_*e*_ = 29*μm*, which, from our derivations, denotes the height of the region at the colony edge where the volume doubles during one cellular doubling time. **(B)** To validate the value of *L*_*e*_ we get from (A), we sought to directly determine the biofilm thickness, *L*_0_, that defines the region that doubles in volume during one cellular doubling time, *τ* . To this end, we compared consecutive interferometry images of *V. cholerae WT* grown on 1.5% agar, as described in this cartoon. We measured the edge volume for different heights *L*_0_ at each initial time point, *V*_*initial*_. These edge volumes include all volume for which *a > x*_1_, where *x*_1_ is the value of the radius where biofilm thickness is *L*_0_. We then measured the edge volume at the second time point, *V*_*final*_, a time *δt* later, where again *a > x*_1_. **(C)** We plotted the ratio of final to initial edge volume against different *L*_0_ values (the purple line). We also plotted the expected *V*_*final*_*/V*_*initial*_ ratio, which is 2^*δt/τ*^ (the green line). Note, the mean time between consecutive measurements, 0.65 hr, is a little less than the measured mean cellular doubling time (*τ* ) of 0.82 hr, ; thus, during the time that elapsed an exponentially growing region should increase its volume by a factor of *∼* 1.7. The intersection point of these two lines thus represents the thickness, *L*_*e*_, where the volume doubles during one cellular doubling time (i.e., *τ* ). We observe that *L*_*e*_ estimated from measurements of seven different strains and conditions in (A) aligns with the direct, empirical measurement of *L*_*e*_. The shaded region for the volume ratio represents the standard deviation from four consecutive time points for one replicate, and we propagated the errors in *τ* and *δt* to determine the standard deviation for 2^*δt/τ*^ . **(D)** Using the empirically measured values of *L*_*e*_, we predicted (Δ*a/*Δ*t*) with Equation 5. The bar plot compares the predicted (Δ*a/*Δ*t*) to the mean measured (Δ*a/*Δ*t*). Our predictions closely match the measured rates, with a mean absolute error of 2*μm/hr*.

To validate our estimate of *L*_*e*_, we used an empirical approach. Note that in our derivation we define the SCNR-region of interest based on its maximum height, *L*_0_. We then define the specific SCNR that doubles in volume during one cellular doubling time by its maximum height, which we now term *L*_*e*_. We thus measured biofilm’s edge volume at two sequential time intervals separated by *δt*. We calculated the edge volume of the biofilm at a specific edge height ( *L*_0_) for the first time point (Fig. 5B). Then, using the same distance from the center for that specific *L*_0_, we calculated the volume in this now larger region in the second time point after a time *δt*. As we separately measured the cellular doubling time, we then compare the ratios of the final to initial volume to 2^*δt/τ*^ . The *L*_*e*_ value that gives *V*_2_*/V*_1_ = 2^*δt/τ*^ is the *L*_*e*_ value to which the volume of the biofilm edge doubles (Fig. 5C). We performed these measurements for *Vibrio cholerae WT* grown on 1.5%, 3%, and 5 % agar. We find the mean *L*_*e*_ to be 33 ± 1, 24 ± 0.5, 29 ± 0.25 *μm* for 1.5%, 3%, and 5 % agar, respectively. Here the errors represent the standard deviation of *L*_*e*_ between means of three different replicates. Thus, our estimated measurement value of *L*_*e*_, from seven diverse strains and species, indeed aligns with the *L*_*e*_ value that corresponds to the volume of the edge that doubles.

### F. Predicting range expansion rate from *θ, L*_*e*_, and *τ*

We now have all of the information needed to predict the range expansion rate. Focusing just on the volume at the edge that doubles, we have

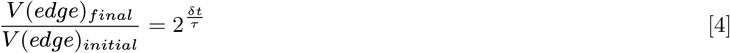

Using volume expression of the edge from Equation 2, we can derive an exact equation for the change in *δa* during time *δt* (See SI Fig. S11 and Eqn.S27):

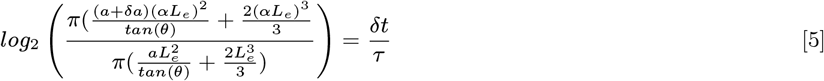

We numerically calculated the range expansion rate using this equation for biofilms grown on three different agar percentages. We find that predicted range expansion rates are very similar to experimentally measured values (Fig.5 D). In fact, the average absolute error is 2 *μm* /hr (See SI for dependence of predictions with *L*_*e*_ and *a*).

Finally, we note that Eqn.3 suggests that a strain with a long doubling time can have a faster *da/dt* than a strain with a shorter doubling time, so long as its *θ* is small enough. We thus use sensitivity analysis to determine which variable *τ* or *θ, da/dt* is more sensitive to (25). The sensitivity of *da/dt* with *τ* and *θ* is given by:

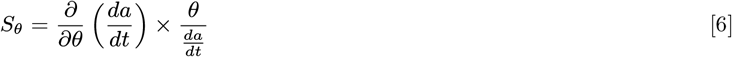

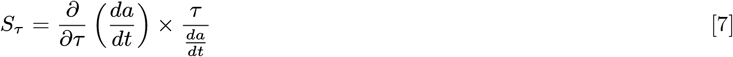

Thus, the ratio of two of these sensitivities is given by

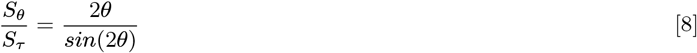

We first see that this ratio is independent of the value in *τ* . This suggests that the sensitivity of *da/dt* does not depend on the particular value of *τ* . Crucially, we also note that for 0 *< θ <* 90°, this ratio is always greater than 1. Thus, *da/dt* is functional is more sensitive to changes in *θ* than changes in *τ*, suggesting that the range expansion rate is more sensitive to the geometry of a biofilm than to the cellular doubling time.

## 3. Discussion

Here, we showed that geometry strongly governs the range expansion rate of growing biofilms. We used white light interferometry to measure monoclonal colony topographies, and found that nascent colonies exhibit spherical cap shapes. Eventually, the colony center is too tall to continue growing exponentially, and the colony as a whole no longer resembles a spherical cap. However, we found that the edge continues to resemble a spherical cap, and is best described as a spherical-cap-napkin-ring (SCNR). We developed analytical expressions for this geometry, and showed that with three parameters (two of which are geometrical) we can predict range expansion rates with a high degree of accuracy. We show that, to leading order, the range expansion rate exhibits a very strong correlation with 1*/*(*τtan*(*θ*)). Finally, we demonstrated that the range expansion is more sensitive to changes in *θ* than it is to changes in *τ* . Thus, biofilm growth strongly depends on biofilm geometry.

It is important to note that the contact angle at the colony’s edge may vary based on the surface conditions where the colony is growing. For instance, changes in the environmental nutrient concentration or surface properties during colony growth could potentially lead to changes in the contact angle. Thus, the constant contact angle we observe is likely due to the consistency of agar experiments in the lab, and not a general property of biofilms. However, the relationship between contact angle and range expansion rate, and in particular Eqn. 3, should hold even as contact angle changes. Future work may investigate dynamic changes to the contact angle and range expansion rate.

Recent works have suggested that shape and morphology of expanding biofilms are impacted by the balance between two forces, friction-like interactions between bacteria and the substrate, as well as surface tension-like interactions between bacteria (8, 15, 18, 22, 26, 27). This idea is consistent with our observation that short biofilms exhibit spherical cap shapes, much like liquid droplets. For sessile liquid drops, the spherical cap shape arises as the balance between interfacial tensions sets the contact angle, and surface tension then minimizes surface area, producing a spherical cap. Further, many references model increasing agar percentage as increasing friction (15, 28, 29); we observe that the contact angle increases with increasing agar percentage, as this idea would suggest. Thus, our experimental observations provide further evidence consistent with this force-based interpretation of biofilm morphology. However, further work is necessary to directly demonstrate that the contact angle and morphology can be directly related back to these “macroscopic” forces.

Surprisingly, a consistent contact angle is observed at the biofilm edge even for strains that exhibit wrinkling, as seen in our study with a high EPS-producing strain (*V. cholerae EPS+*). Intriguingly, the wrinkled colony falls on the same trend line as smooth colonies. This consistency suggests that the spherical cap napkin ring analysis remains a valid and precise method for predicting the range expansion rate of wrinkled colonies. Future work may test if the mechanical changes that lead to extreme buckling and delamination modify the contact angle and thus have a commensurate impact on the range expansion rate (22, 30, 31).

Note that while we find that the spherical cap and spherical cap napkin ring are the most accurate geometrical descriptions of nascent and mature biofilms, the cone and cone napkin ring should still produce a range expansion rate that is proportional, to leading order, to ∝ *L*_*e*_*/*(*τtan*(*θ*)), and thus should still produce relatively accurate predictions. In fact, so long as a model includes the idea that range expansion rate should be inversely proportional to *tan*(*θ*), it is likely to be relatively accurate.

The geometry of the propagating front of a growing colony may also have ecological and evolutionary consequences. In this work we have only investigated monoclonal colonies. For future work, it would be interesting to study the competition between different strains in a mixed species biofilm, and look at the consequence of different contact angles at the propagation front. Moreover, cellular populations can assume a variety of shapes from smooth to rough surfaces and branching morphologies. Structural variation emerges from, e.g., demographic noise and growth instabilities (32–34). While these effects already have complex and profound consequences, future works may investigate how those effects combine with, and compare to, the effects of biofilm geometry.

Finally, our results suggest that to understand biofilm fitness, cellular doubling time and biofilm contact angle must both be measured. Thus, to understand the fitness of biofilms in real environments, the contact angle must be measured on the actual surfaces on which they grow (or surfaces with similar material properties). Thus, biofilm and substrate material properties must be considered for their effect on contact angle, just as environmental nutrient concentrations are considered for their effect on cellular doubling times.

## 4. Methods

### A. Biofilm growth on agar plates

Cell stocks (See SI Table 1 for strains used) from 80° C were cultured overnight at 37° C in a shaking incubator set at 220 rpm in sterile liquid LB medium prepared using 10g tryptone, 5g yeast extract, and 10g NaCl in 1L of water (35). These stocks were subsequently diluted in LB to an optical density (OD_600_) of 1. To facilitate growth from individual cells, these OD 1 cultures were further diluted in LB by factors ranging from 10^6^ to 10^7^. 3 *μl* of the diluted samples were then inoculated on LB agar plates, which were prepared by sterilizing liquid LB medium to which was added the required agar concentration (1.5, 3 and 5 %w/v) for different measurements. For time-lapse experiments, the plates were incubated at 23° C for roughly 15 hours prior to their placement under the interferometer for measurements. Plates were incubated at 23° C, except during the periods they were positioned under the interferometer, which operated at room temperature (*≈* 23° C).

### B. Image acquisition and analysis

We imaged grown colonies using Zygo Zegage Pro interferometer with a 50X Mirau objective. To prevent evaporation of the agar plates during time lapse measurements, we placed a plastic casing around the interferometer. This prevented drying of the plates and the media, and thus yielded consistent measurements. Similar methodologies have been previously used in Bravo, et al., (7).

Initially, we examined the colony morphology to ensure that we considered only colonies originating from individual cells. Colonies resulting from the merger of two cells were discernible due to their “figure-8” shape. Given the symmetry of colonies grown from single cells, we imaged only a quarter of larger biofilms to reduce acquisition time.

All the images were processed using a custom python script available in GitHub. To subtract the agar background from the biofilm topography, we computed a best-fit polynomial based on the image regions without the colony and subtracted this from the entire image. This step negated any measurement artifacts arising from the inclination of the agar. We tried three distinct thresholds (5, 10 and 20 %) of colony height to select the colony’s outer edge and subsequently fit an best fit circle and extract the center of biofilm (refer to Supplementary Information). For contact angle measurements, we extracted 20 different profiles around the colony originating from center of the colony to the outer edge. The contact angle was quantified as the mean of the maximum angle observed along these profiles.

### C. Doubling time measurements

Cell stocks from a − 80°C freezer were cultured overnight in liquid LB medium and then diluted to an optical density (OD_600_) of 1. This culture was further diluted by a factor of approximately 10^4^ in liquid LB. To ensure the cells remained in the exponential growth phase, the diluted sample was incubated at 37°*C* in a shaking incubator for ~ 3 hours. Subsequently, this sample was spread on agar plates for measurements. Imaging was performed using a Nikon Confocal Microscope equipped with a 20X objective, over an approximate duration of 6 hours at 23°*C*. The dilution process ensured a sufficient number of individual cells within the microscope’s field of view, allowing clear observation of cell-doubling events. Analysis of these doubling events was conducted using the Fiji-Image J software. Given the clear visibility of doubling events, we recorded visually the time taken for one cell to fully separate and double into two daughter cells.

## ACKNOWLEDGMENTS

P.J.Y. acknowledges funding from the NIH National Institute of General Medical Sciences (grant no. 1R35GM138354-01) and NSF Biomaterials (grant no. BMAT2003721). B.K.H. acknowledges funding from NSF Biomaterials (grant no. BMAT2003721).

