## SupplementText for "The biophysical basis of bacterial colony growth"

### 1. Analysis of biofilm shape goodness of fit

**A. Comparing experimental topographies to different shapes.** We first experimentally measured different parameters required for calculating the theoretical shape. These parameters included the centroid coordinates $(x_c, y_c)$, the radius $a$, and the height $h$ of the colony. Subsequently, for each set of planar coordinates $(x, y)$ from the actual biofilm topography, we computed the corresponding $z$ values using the appropriate shape equations. The disparity between the predicted $z$ values and the actual $z$ values derived from experimental topographies was utilized to determine the residual. Below, we provide descriptions of the methods employed to calculate different parameters for various shapes.

***A.1. Spherical cap.*** A spherical cap is defined by the equation of a sphere

$$(x - x_c)^2 + (y - y_c)^2 + (z - z_c)^2 = (R)^2 \tag{S1}$$

which includes three centers $(x_c, y_c, z_c)$ and the radius $(R)$. We measured $x_c$ and $y_c$ by finding the coordinates with maximum height $(h)$ in a biofilm topography (See Methods section for details on the center calculations). The radius $(a)$ is measured from the center to the edge of a topography. The $z_c$ for spherical cap shape, when the origin of the coordinate system is at the center of the sphere, is given by the difference between the radius of the sphere (R) and the height (h).

***A.2. Gaussian.*** A Gaussian function with a two-dimensional domain is defined as

$$z = A \exp\left(\frac{(x - x_c)^2}{2\sigma_x^2} + \frac{(y - y_c)^2}{2\sigma_y^2}\right) + C \tag{S2}$$

Here, $A$ represents the amplitude of the Gaussian, while $x_c$ and $y_c$ denote the respective centers of the shape. C represents the offset of the Gaussian denoting the offset in the position of z coordinates. The parameters $\sigma_x$ and $\sigma_y$ indicate the spreads of the Gaussian function in the $x$ and $y$ directions. We used height of the biofilm, $h$, for the amplitude of the Gaussian $(A)$ with zero offset. Similar to the calculation for a spherical cap, we identified the coordinates with the maximum height in the topography to measure $x_c$ and $y_c$ for the Gaussian. $\sigma_x$ and $\sigma_y$ were calculated by measuring the full-width-half-maximum (FWHM) in both the $x$ and $y$ directions and equating it with $2\sqrt{2ln2}\sigma$(1).

***A.3. Cone.*** The equation for a cone in Cartesian coordinates is given by

$$\frac{(x - x_c)^2 + (y - y_c)^2}{c^2} = (z - h)^2 \tag{S3}$$

where $c$ represents the ratio of the radius $(a)$ to the height $(h)$ of a cone. We used similar approach as described above to calculate centers, radius and height of biofilm, i.e., we measured the maximum point height of the biofilm to get $h$, and the respective (x,y) coordinates as center of the topography.

***A.4. $R^2$ values.*** Figure S1 shows the $R^2$ values obtained from comparing the experimental topographies with the above mentioned approach and their respective perfect shapes. The spherical cap always has the highest $R^2$ value, and for all strains studied ($R^2 > 0.95$).

**B. Least squares fit.**

***B.1. All free parameters.*** To further investigate the differences between two closely related shapes, the spherical cap and Gaussian, we applied a least squares fitting algorithm to all measured topographies. Initially we allowed all parameters to vary, enabling us to compare the best fit values to the actual measured values.

For the spherical cap, we discovered excellent agreement between the best fit and measured values for the radii and center coordinates (Fig. S2). It's important to note that in this context, the radius refers to the radius of the corresponding sphere $(R)$ rather than the in-plane radius of the spherical cap $(a)$.

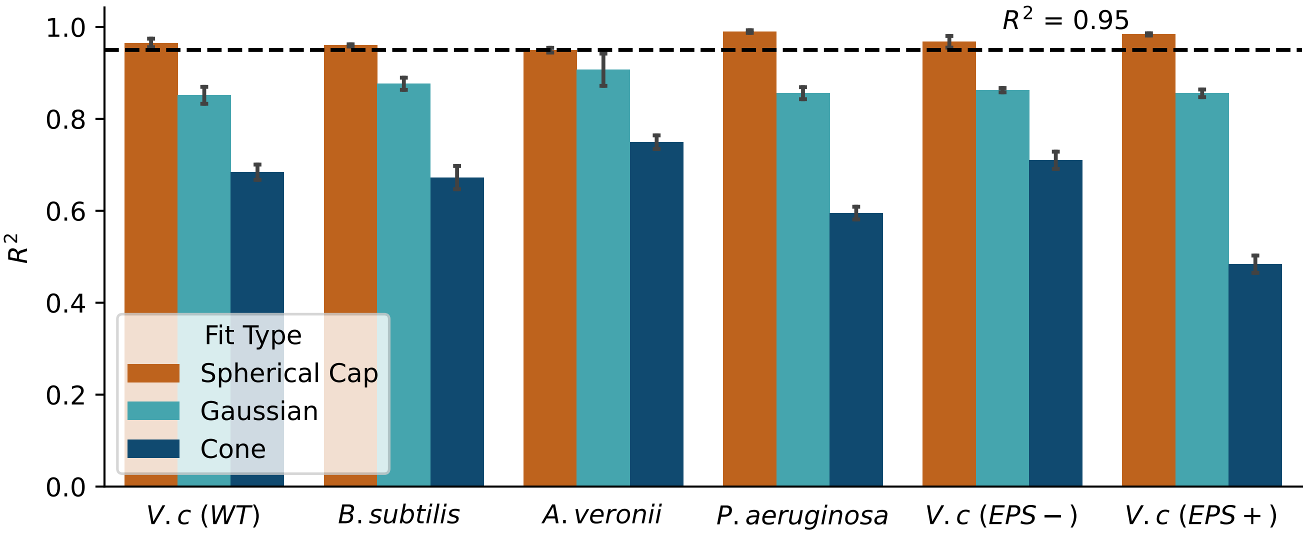


**Fig. S1.** The $R^2$ value, obtained from comparing the z coordinates obtained from different shape equations to the measured topography, is plotted for six different strains. Error bars represent the standard deviation from three different biofilms grown on same agar plate. The colonies were grown on 1.5% agar plates for approximately 24 hours at $23°$C, with the exception of *B. subtilis*, which was grown for approximately 96 hours to attain a similar size as the other strains. For all strains measured, we observe $R^2 > 0.95$ for spherical cap shape.

In contrast to the spherical cap, for the Gaussian shape, we found that the fitted values deviated significantly from the measured values for all parameters except the centers. In particular, the amplitude obtained from the fitting was much larger than the measured amplitude values (the height of the biofilm) (Fig. S3). This suggests that the only way to achieve a good least square fit for the Gaussian is to fit a Gaussian with an artificially large amplitude, effectively using only the peak of the Gaussian to match the topography.

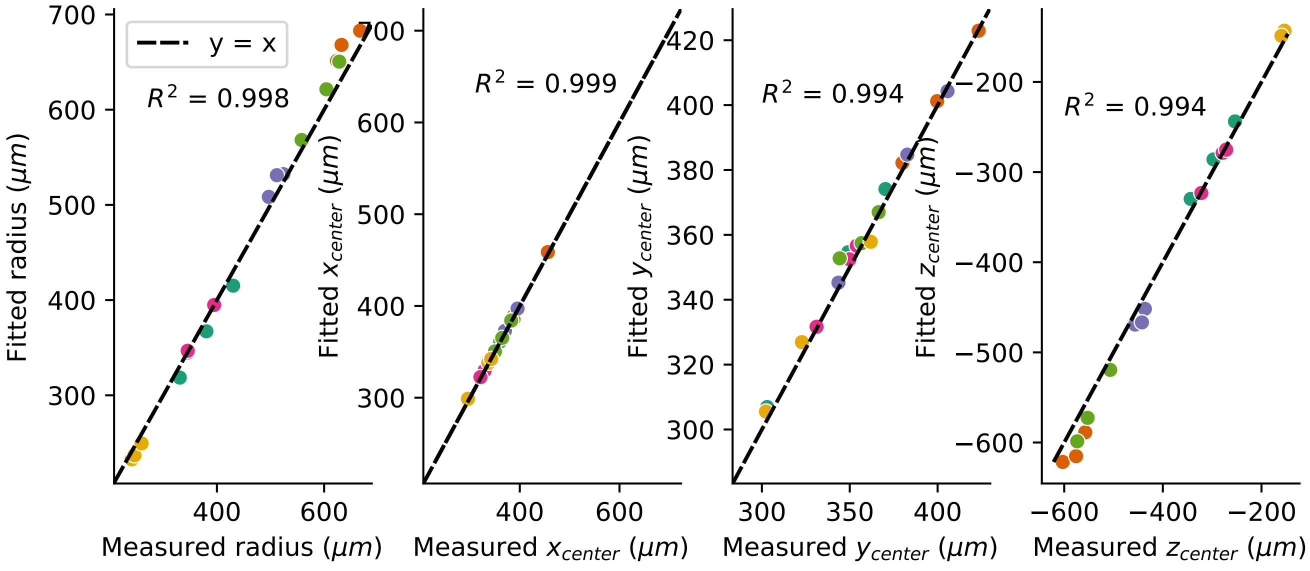


**Fig. S2.** The least square fit parameters from fitting a spherical cap shape are plotted against the corresponding measured parameters for all six strains. Different colors in scatter plot represent different strains (as defined in Fig. 1 of the main text). We see an excellent agreement ($R^2 > 0.98$) between fit values and measured values for the spherical cap shape.

***B.2. Few free parameters.*** We next fixed some of the parameters required for fitting to see how the residual compares between spherical cap and Gaussian shapes. For the spherical cap, we fixed the $x_{center}$ and $y_{center}$ of biofilm and only allowed the

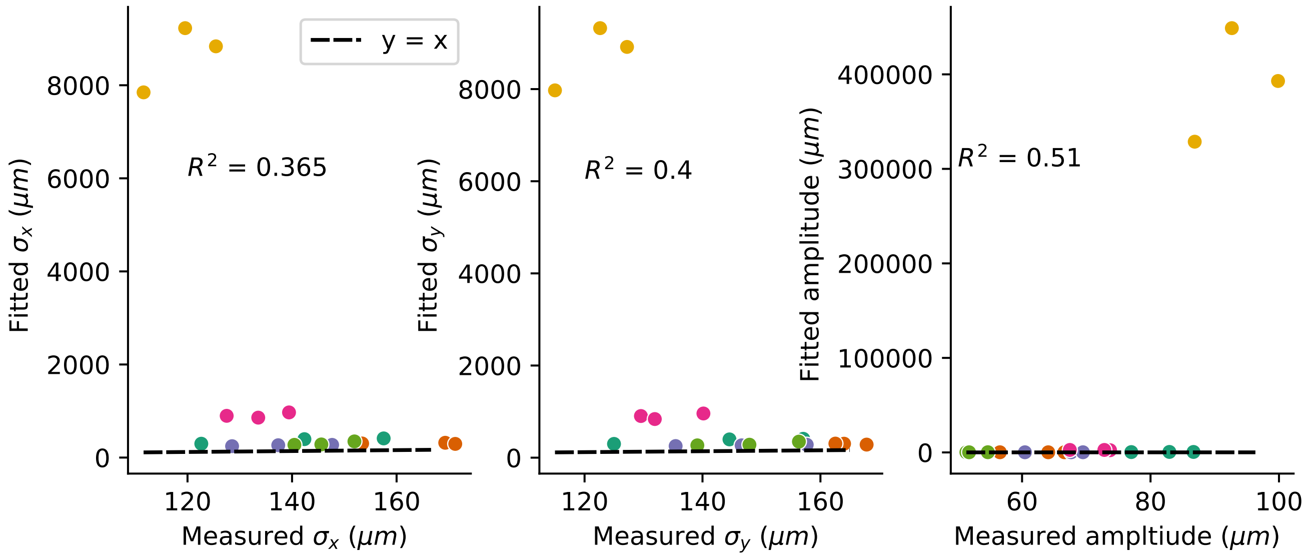


**Fig. S3.** The least squares fit parameters from fitting Gaussian shape are plotted against the measured values for all six strains. Different colors in the scatter plot represent different strains (as defined in Fig. 1 of the main text). We observe a poor agreement between the fitted values and the measured values for the Gaussian least squares fit. Note that, even when we calculate $R^2$ without the outliers, which in this case originate from *V. cholerae (EPS+)*, we still obtain $R^2 < 0.55$ for the above fitted variables. The fitted value for centers are not shown in the above plots. The least square fit provides a good prediction for only centers of the Gaussian ($R^2 > 0.99$).

$z_{center}$ and radius ($R$) to vary. For fitting the Gaussian, we again fixed $x_{center}$ and $y_{center}$, along with the offset, so that the algorithm doesn't fit an artificially large amplitude. The residual from the Gaussian fit is much larger than the residual from the spherical cap fit for all strains (Fig. S4).

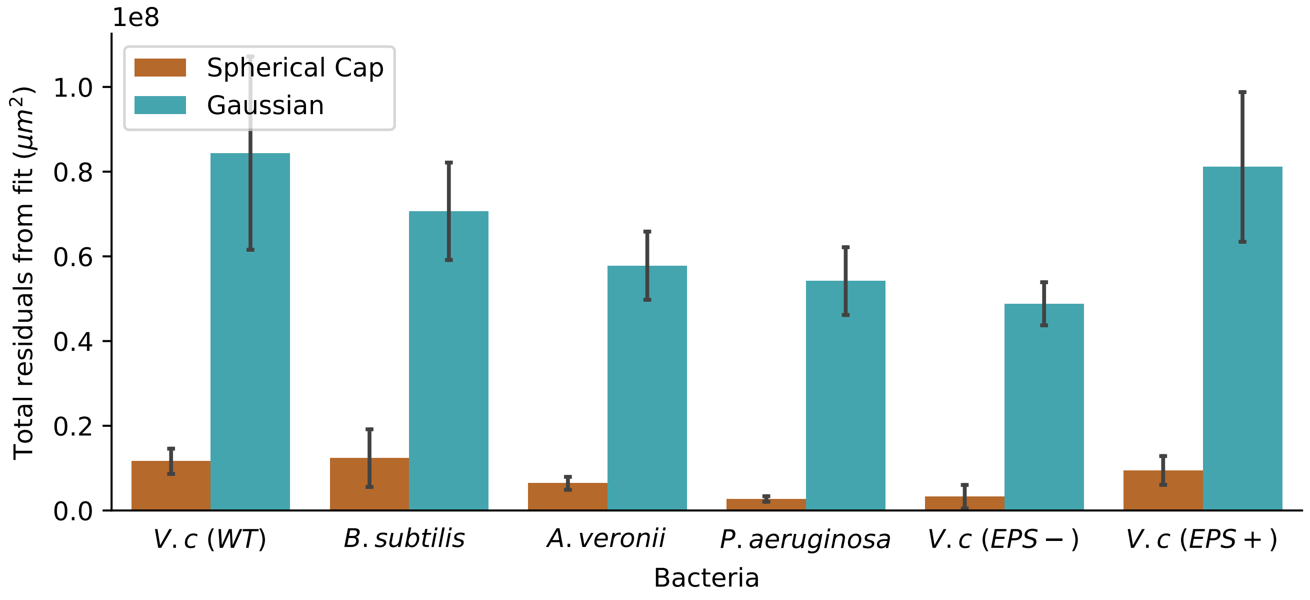


**Fig. S4.** The total residuals obtained from the difference between measured values and those obtained from the least squares fits are plotted for two different shapes. In both cases, the centers were kept fixed, and an offset constraint was imposed on the Gaussian shape. An offset constraint was used to prevent the fitting algorithm from fitting an inappropriately high-amplitude Gaussian. Despite this constraint, the Gaussian fit yields higher residuals compared to the fit using a spherical cap shape. This result emphasizes the superior fit of the spherical cap shape even in such a constrained scenario.

**C. Volume comparison.** As a final step in assessing the goodness of fit of the spherical cap, we measured the volume of each of the topographies and compared it to the volume of a perfect spherical cap with the same radius ($a$) and height ($h$). Our findings indicate that the measured volumes and the volumes of the perfect spherical caps exhibit excellent agreement ($R^2 = 0.96$) (Fig. S5).

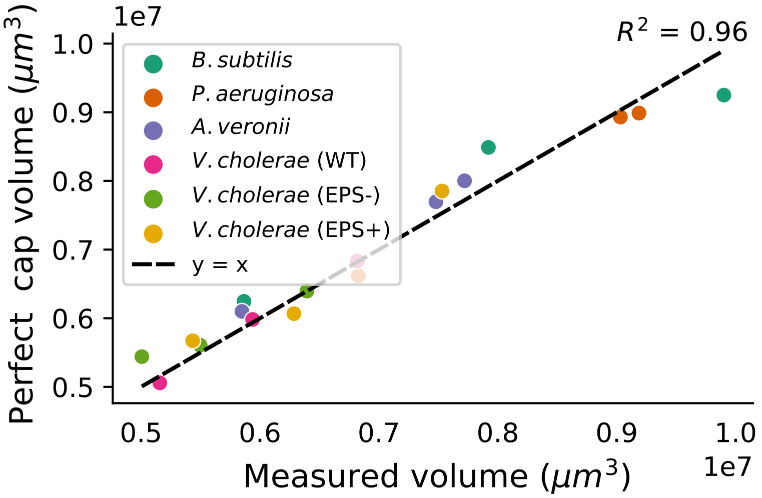


**Fig. S5.** The exact spherical cap volume for the measured radius and height is plotted against the measured volume of the biofilm topography. An $R^2$ value of 0.96 indicates that these colony volumes are well described by the spherical cap shape.

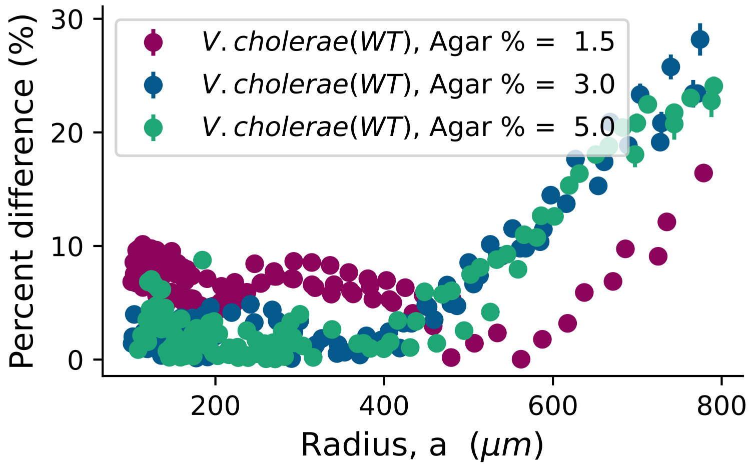


**Fig. S6.** The percentage difference between the measured volume and the volume of a perfect spherical cap with measured $a$ and $\theta$ is plotted against $a$. Different colors represents data from different agar percentages (as indicated in the inset); three different biofilm replicates were used for each condition. Across all different agar percentages, we observed that the percentage difference begins to increase noticeably after $a$ exceeds approximately 500 $\mu m$. The error bars in the percentage difference (which, in some cases, are difficult to discern due to the size of the scatter plot) represent errors propagated from the measurements of $a$ and $\theta$.

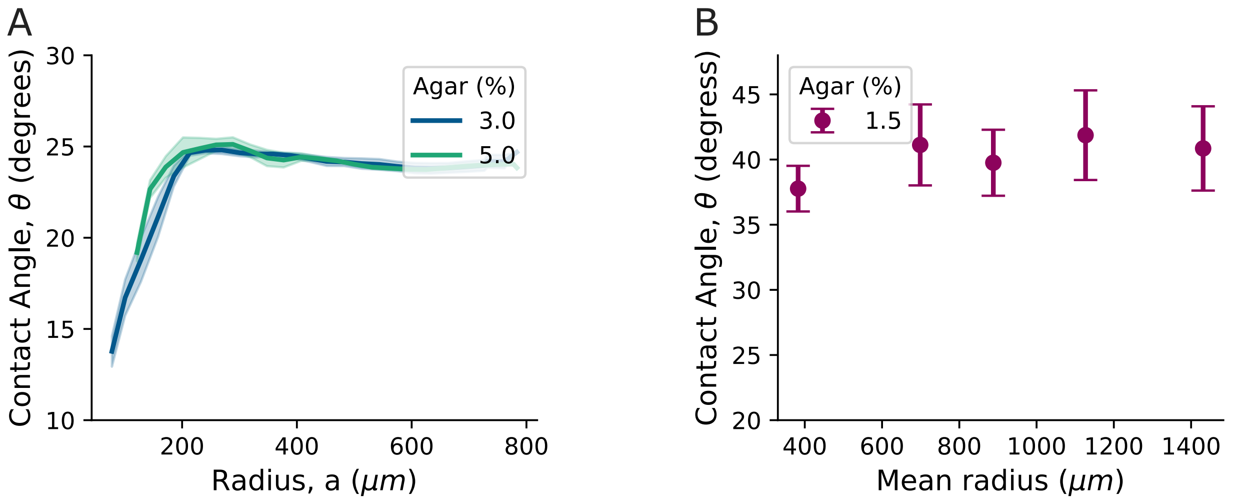


**Fig. S7. A)** The contact angle, $\theta$, is plotted against the colony radius for *V. cholerae (WT)* grown on 3% and 5% agar plates. The solid lines represent the LOESS-averaged values, and the shaded region represents the standard deviation from three replicates. **B)** In a separate measurement, we observed the growth of colonies of *V. cholerae (WT)* over a period of approximately 3.5 days, and measured $\theta$. This long-term measurement demonstrates that the contact angle remains constant even over extended time scales. Note that, for this experiment, agar plates were inside an incubator except during the time of measurement, resulting in a different contact angle compared to the conditions shown in the main text, where the plates were kept under the interferometer inside a plastic casing for the duration of the measurements.

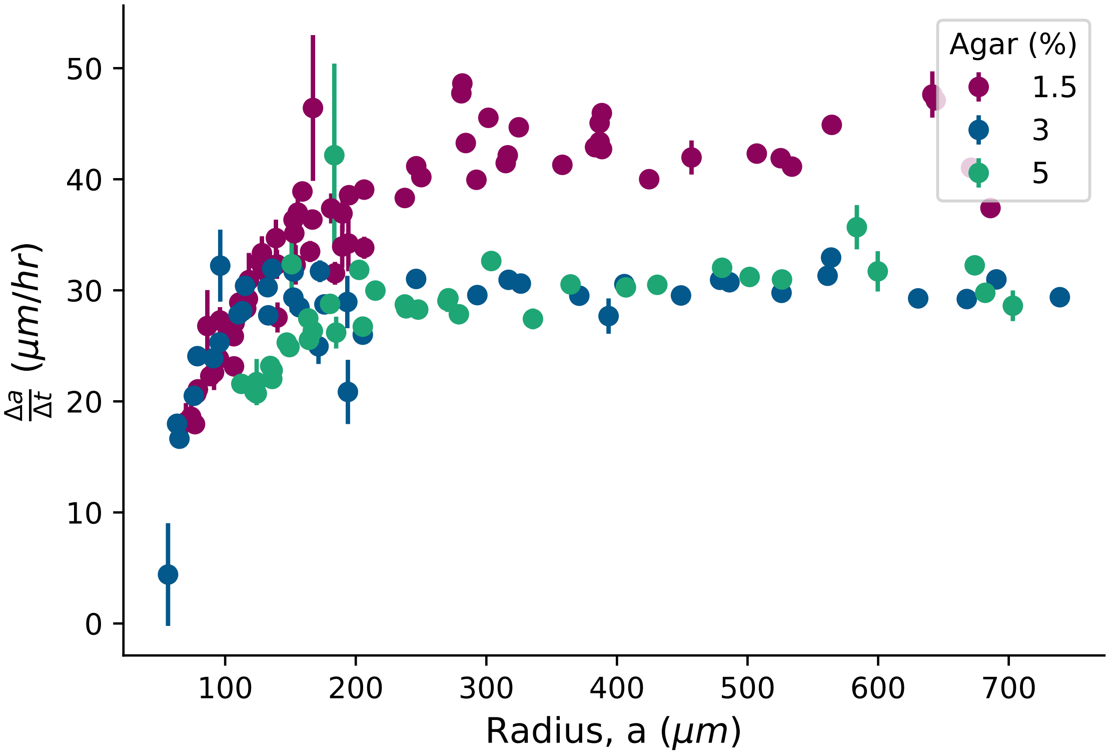


**Fig. S8.** The range expansion rate for *V. cholerae (WT)* colonies grown on different agar percentages is plotted against the colony radius. The range expansion rate is determined by finding the slope of every three consecutive measurements of $a$ versus time, with error bars representing the errors obtained from the slope fit of linear regression. We observe that after an initial increase in $\Delta a/\Delta t$, the range expansion rate saturates.

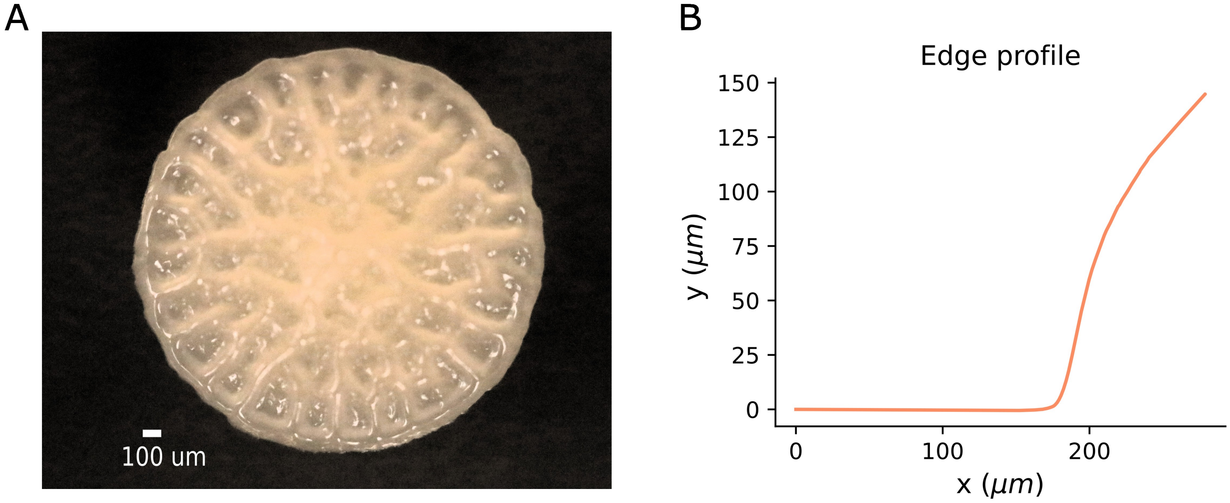


**Fig. S9.** *V.cholerae (EPS+)* is grown from a single cell at $23^{\circ}$ C for approximately 170 hours. Over this period, the colony exhibits wrinkling and buckling at its center. Panel **A** displays an image of the entire colony captured using a Canon MP-E 65mm camera. Subsequently, we also measured the topography of the same colony to examine its edge. Panel **B** demonstrates that, even in the presence of wrinkle structures within the colony's center, the colony's edge retains a finite contact angle.

### 2. Mathematical derivations

#### A. Geometry of the spherical cap napkin ring.

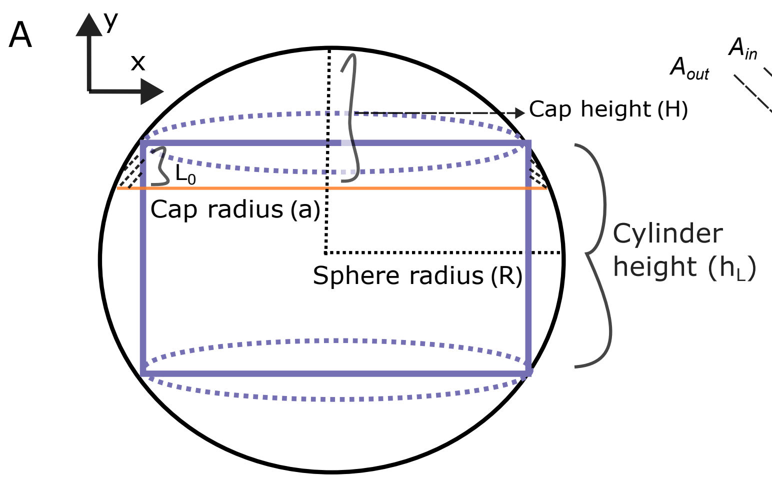


**Fig. S10.** These cartoons define how we describe spherical-cap-napkin-rings. **A)** A cylinder with a height of $h_L$ is subtracted from a sphere, with both the sphere and cylinder sharing the same center. In our analysis, we concentrate on the remaining volume within the spherical cap (the crosshatched region) rather than considering the entire sphere. **B)** In our derivation, we subtract the surface areas $A_{\mathrm{out}}$ and $A_{\mathrm{in}}$ and integrate over the height $L_0$ to obtain the volume of the spherical cap napkin ring (SCNR).

***A.1. Spherical cap napkin ring volume.*** Here we derive the volume of a spherical cap napkin ring (SCNR). We first consider a cylinder of height $h_L$ which is carved out from a sphere with a radius $R$. While the derivation of the leftover volume from the entire sphere is well-known and referred to as the napkin ring volume, our objective is to determine the leftover volume within the spherical cap region (as shown in cross-hatched region in Fig. S10), characterized by the edge height ($L_0$). This volume, the SCNR volume, is obtained through the integration of horizontal cross-sectional areas over the height of the spherical cap napkin ring, as described by Equation S4.

$$V_{nap} = \int_{y_1}^{y_2} dy(A_{out} - A_{in}) = \int_{y_1}^{y_2} dy(\pi R_{out}^2 - \pi R_{in}^2) \quad \text{[S4]}$$

Here, $A_{out}$ and $A_{in}$ refer to the circular areas at the same $y$ value that extend all the way to the outer and inner edges of the SCNR ($A_{out}$ and $A_{in}$, respectively); $R_{\mathrm{out}}$ and $R_{\mathrm{in}}$ thus refer to the radial distances from the center of the spherical cap to the outer edge of the sphere and to the point where the edge height line intersects the base of the spherical cap, respectively, which are geometrically defined as:

$$R_{in} = \sqrt{R^2 - \left(\frac{h_L}{2}\right)^2} \quad \text{[S5]}$$

$$R_{out} = \sqrt{R^2 - y^2} \quad \text{[S6]}$$

With simple geometry, we obtain $y_1 = R - H$, $y_2 = R - H + L_0$. Thus,

$$V_{nap} = \pi \int_{R-H}^{R-H+L_0} dy(\left(\frac{h_L}{2}\right)^2 - y^2) \quad \text{[S7]}$$

$$\text{[S8]}$$

The spherical cap geometry allows us to express $R - H = \frac{a}{\tan(\theta)}$ and $\frac{h_L}{2} = \frac{a}{\tan(\theta)} + L_0$, where $\theta$ is the contact angle between the edge of the spherical cap and the horizontal. Upon making these substitutions, we arrive at the final expression for the volume of the SCNR as a function of $a$, $\theta$, and $L_0$.

$$V_{nap} = \pi L_0^2 \left(\frac{a}{\tan(\theta)} + \frac{2L_0}{3}\right) \quad \text{[S9]}$$

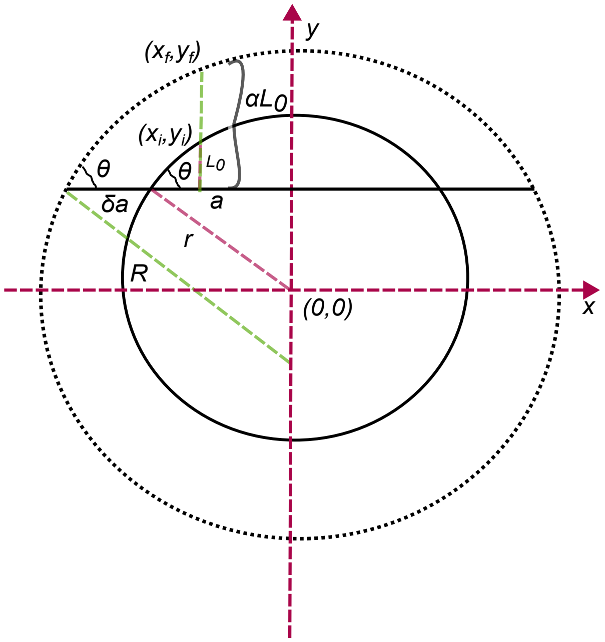


**Fig. S11.** We consider a spherical cap with radius $a$ that undergoes expansion, maintaining a constant angle $\theta$, to reach a new radius $a + \delta a$. Simultaneously, the edge height grows from $L_0$ to $\alpha L_0$. The encompassing sphere's radii change from $r$ to $R$. Our goal is to calculate the rate of range expansion, $\frac{\delta a}{\delta t}$. This cartoon also defines $(x_i, y_i)$ and $(x_f, y_f)$.

### B. Derivation of spherical cap napkin ring range expansion rate.

***B.1. Relationship between vertical growth and $\delta a$ for SCNR.*** To create an analytical prediction for the range expansion rate of a SCNR, we first must determine how much the height of a SCNR increases when its radius increases a small amount. We thus define a couple of geometric parameters, as shown in Fig. S11; $\alpha$ represent the change in edge height as the spherical cap expands by $\delta$a. We now solve for $\alpha$ using SCNR and spherical cap geometry.

The equation for a circle defines a vertical cross section of the sphere of the spherical cap. Setting the origin at the center of the initial circle, the equations for these circles are then as follows, where $r$ and $R$ are the radii of the initial and final spheres and $y_c$ is center of the larger sphere. Let $(x_i,y_i)$ and $(x_f,y_f)$ denote the points on the initial spherical cap at edge height $L_0$ and the points on the final spherical cap at edge height $\alpha L_0$, respectively (see Fig. S11). These points, as they reside on the surface of the spherical cap, satisfy the following equation of a circle.

$$x_i^2 + y_i^2 = r^2 \tag{S10}$$
$$x_f^2 + (y_f - y_c)^2 = R^2 \tag{S11}$$

Since both points lie on same y-z plane, we have $x_i = x_f$. $y_i$, $y_f$, $r$ and $R$ can be simply determined by using spherical cap geometry .

$$y_i = \frac{a}{\tan(\theta)} + L_0 \tag{S12}$$
$$y_f = \frac{a}{\tan(\theta)} + \alpha L_0 \tag{S13}$$
$$r = \frac{a}{\sin(\theta)} \tag{S14}$$
$$R = \frac{a + \delta a}{\sin(\theta)} \tag{S15}$$

To find the downward shift of the center of the larger sphere of radius $R$, we find the difference between the vertical side lengths of two similar triangles

$$|\mathrm{y_c}| = \frac{a + \delta a}{tan(\theta)} - \frac{a}{tan(\theta)} \tag{S16}$$
$$|\mathrm{y_c}| = \delta a cot(\theta) \tag{S17}$$

We can now rewrite equation S11 in terms of S10 as

$$x_i^2 + (y_f - y_c)^2 = R^2 \quad \text{[S18]}$$
$$(r^2 - y_i^2) + (y_f - y_c)^2 = R^2 \quad \text{[S19]}$$

Substituting all the values of equations S12, S13, S14, S15, S17 in S19,

$$\left(\frac{\mathrm{a}}{\sin(\theta)}\right)^2 - \left(\frac{a}{tan(\theta)} + L_0\right)^2 + \left(\frac{a}{tan(\theta)} + \alpha L_0 - (-\delta a cot(\theta))\right)^2 = \left(\frac{a+\delta a}{sin(\theta)}\right)^2 \quad \text{[S20]}$$

[S21]

If we solve for $\alpha$ and then simplify, we get

$$\alpha = \frac{\sqrt{\left(\frac{a+\delta a}{\sin(\theta)}\right)^2 - \left(\frac{a}{\sin(\theta)}\right)^2 + \left(\frac{a}{\tan(\theta)} + L_0\right)^2} - \frac{\delta a}{\tan(\theta)} - \frac{a}{\tan(\theta)}}{L_0} \quad \text{[S22]}$$
$$\alpha = \frac{1}{L_0}\left(\sqrt{\csc^2(\theta)\left(2a\delta a + \delta a^2\right) + (L_0 + a\cot(\theta))^2} - (a+\delta a)\cot(\theta)\right) \quad \text{[S23]}$$

***B.2. Analytical expressions of exact and approximate SCNR range expansion rates.*** The volume before and after the SCNR radius expansion described in the previous section can be written as the following equations.

$$V_{initial} = \frac{\pi L_0^2}{\tan(\theta)}\left(\frac{a}{\tan(\theta)} + \frac{2L_o}{3}\right) \quad \text{[S24]}$$
$$V_{final} = \frac{\pi\alpha^2 L_0^2}{\tan(\theta)}\left(\frac{a+\delta a}{\tan(\theta)} + \frac{2\alpha L_o}{3}\right) \quad \text{[S25]}$$

If the SCNR volume expansion occurs in time $\delta t$, and the source of this expansion is the exponential growth of cells in the SCNR, then this expansion can be related to the doubling time of the bacteria ($\tau$) as,

$$2^{\frac{\delta t}{\tau}} = \frac{V_{final}}{V_{initial}} = \frac{\alpha^2\left(\frac{a+\delta a}{\tan(\theta)} + \frac{2\alpha L_o}{3}\right)}{\left(\frac{a}{\tan(\theta)} + \frac{2L_o}{3}\right)} \quad \text{[S26]}$$
$$2^{\frac{\delta t}{\tau}} = \alpha^2 \frac{1 + \frac{\delta a}{a} + \frac{2\alpha\tan(\theta)}{3}\frac{L_0}{a}}{1 + \frac{2\tan(\theta)}{3}\frac{L_0}{a}} \quad \text{[S27]}$$

Eqn S27 is exact, but does not appear readily solvable. In the experimental regime of $1° < \theta < 70°$, $\frac{2\tan(\theta)}{3} \lesssim \mathcal{O}(1)$. Typical values of $L_0$ and $a$ result in the approximation that $\frac{L_0}{a} \lesssim 0.1$. We therefore Taylor expand around the second term in the denominator with a maximum error of about 1%.

$$2^{\frac{\delta t}{\tau}} = \alpha^2\left(1 + \frac{\delta a}{a} + \frac{2\alpha\tan(\theta)}{3}\frac{L_0}{a}\right)\left(1 - \frac{2\tan(\theta)}{3}\frac{L_0}{a}\right) \quad \text{[S28]}$$
$$= \alpha^2\left(1 + \frac{\delta a}{a} + (\alpha - 1)\frac{2\tan(\theta)}{3}\frac{L_0}{a} - \alpha\frac{4\tan^2(\theta)}{9}\left(\frac{L_0}{a}\right)^2 - \frac{2\tan(\theta)}{3}\frac{\delta a}{a}\frac{L_0}{a}\right) \quad \text{[S29]}$$

With $\delta a \lesssim L_0 \lesssim \frac{a}{10}$, all terms of $\frac{\delta a}{a}$ and higher order terms of $\frac{L_0}{a}$ will be dropped. Additionally, since $\alpha \sim 1$, we can arrive at an approximate expression for the ratio of the final volume to the initial volume as:

$$2^{\frac{\delta t}{\tau}} \approx \alpha^2 = \left(\frac{1}{L_0}\left(\sqrt{\csc^2(\theta)\left(2a\delta a + \delta a^2\right) + (L_0 + a\cot(\theta))^2} - (a+\delta a)\cot(\theta)\right)\right)^2 \quad \text{[S30]}$$
$$= \left(\frac{a}{L_0}\left(\sqrt{\csc^2(\theta)\left(2\frac{\delta a}{a} + \left(\frac{\delta a}{a}\right)^2\right) + \left(\frac{L_0}{a} + \cot(\theta)\right)^2} - (1 + \frac{\delta a}{a})\cot(\theta)\right)\right)^2 \quad \text{[S31]}$$

Solving for $\frac{\delta a}{\delta t}$ yields the radial expansion of a spherical cap napkin ring. To calculate this, the logarithm is taken of both sides of Eqn S31:

$$\ln(2)\frac{\delta t}{\tau} \approx 2\ln(\alpha) \qquad \text{[S32]}$$

And the RHS is simplified using the laws of logarithms. As previously expressed, $\alpha$ typically takes values close to unity, so an approximation like $\ln(1+x) \approx x$ could be taken, but the functional form of $\alpha$ is too complicated. Instead, $\frac{\delta a}{a}$ is assumed to be small and a Taylor expansion is taken to the first order and then simplified, yielding:

$$\ln(2)\frac{\delta t}{\tau} \quad \approx \quad 2\frac{\delta a}{L_0}\left(\frac{\tan(\theta) - \frac{L_0}{a}}{\frac{L_0}{a}\tan(\theta) + 1}\right) \qquad \text{[S33]}$$

which can be solved for $\frac{da}{dt}$:

$$\frac{\delta a}{\delta t} \approx \frac{L_0 \ln(2)}{2\tau}\left(\frac{\frac{L_0}{a}\tan(\theta) + 1}{\tan(\theta) - \frac{L_0}{a}}\right) \qquad \text{[S34]}$$

In the range where $\frac{L_0}{a} \ll 1$ and $\frac{L_0}{a} \ll \tan\theta$, as observed in our experimentally measured data, we obtain:

$$\frac{\delta a}{\delta t} \approx \frac{L_0 \ln(2)}{2\tau tan(\theta)} \qquad \text{[S35]}$$

### 3. Measurement of $L_e$

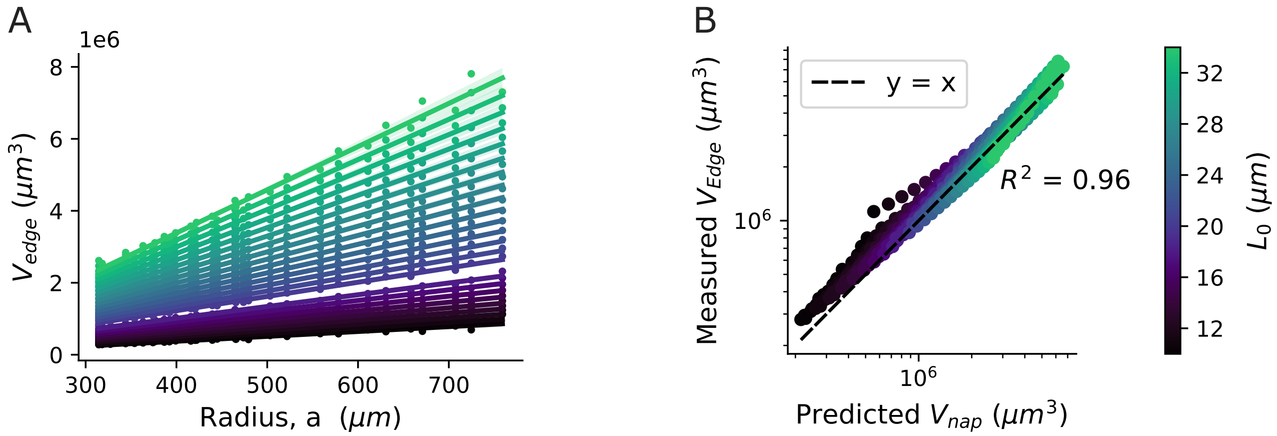


**Fig. S12. A)** The edge volume of a biofilm is plotted against the colony radius, $a$, for different heights $L_0$ in the time regime in which the contact angle is constant. A linear regression line is fitted for each $L_0$ value. For all $L_0$ values, we obtain $R^2 > 0.9$, indicating that the volume of the edge increases linearly with the radius, as predicted by Equation S9. The shaded region in each line represents the 95% confidence interval in the linear fit. **B)** The measured volume at the edge is plotted against the predicted SCNR volume for different $L_0$ values. The resultant $R^2$ value of 0.96 demonstrates excellent agreement between the predicted and measured volumes, suggesting that the spherical cap napkin ring model provides an accurate prediction of the edge geometry.

**A. Error in measurement of $L_e$.** We used the mean doubling time of bacteria as measured from confocal microscopy to determine the edge height at which volume doubles within the measured doubling time (referred to as $L_e$). Thus, error in the measurement of $L_e$ stems from the error in doubling time measurement ($\tau$). For *V. cholerae WT* grown on 1.5% agar, the measured doubling time of bacteria was 49. ± 2. minutes, where the error represents the standard deviation among five different doubling events of distinct bacteria. To investigate the impact of doubling time measurement uncertainty on the measurement of $L_e$ and, ultimately, the prediction of range expansion, we considered two consecutive data points in each replicate to empirically determine $L_e$ across a range of doubling times. We used the mean contact angle for the respective replicates to numerically calculate the range expansion rate using Equation S26. Our results reveal that the prediction of range expansion remains robust, i.e., the uncertainty is relatively low ($\pm\, 2\mu mhr$), even if we consider artificially low doubling times outside the uncertainty range of our measurements (Fig. S13).

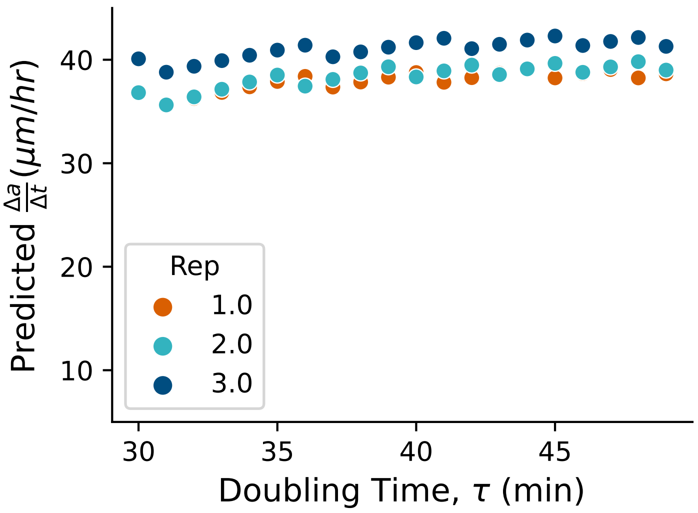


**Fig. S13.** Predicted range expansion rate is plotted against different cellular doubling times for *V.cholerae WT* grown on 1.5% agar. We selected a range of doubling times larger than the error in our doubling time measurements. We then used the mean contact angle and a radius of 800 $\mu$m to calculate the predicted range expansion. We see that predicted range expansion is robust against errors in the measured doubling time even outside the error range.

**B. Dependence of range expansion rate on contact angle ($\theta$) and radius (a) .** Equation S34 suggests that the range expansion rate ($\Delta a/\Delta t$) depends on the contact angle ($\theta$) and also the radius ($a$), while our leading order approximation (Eqn. S35) does not depend on ($a$). To investigate this dependence, we varied the contact angle and predicted the range expansion rate for three different agar percentages. We used the mean $L_e$ measured from three different replicates, the mean doubling time ($\tau$), and a fixed radius ($a = 500\,\mu m$). As discussed in the main text, we observed a strong dependence of the range expansion rate on the contact angle (Fig. S14A).

We then used the mean contact angle ($\theta_{avg}$), mean $L_e$ measured from three different replicates, and the mean doubling time ($\tau$) to explore the dependence of $\Delta a/\Delta t$ with respect to the radius of the biofilm ($a$). Figure S14 B shows, after the contact angle saturates ($a > 300\,\mu m$), the predicted range expansion rate is fairly constant. Given a constant contact angle, as the radius $a$ increases, the fraction $\frac{L_0}{a}$ becomes smaller, and eventually, as approximated in Equation S35, the range expansion rate thus saturates to a constant value. In our particular case, the predicted change in range expansion over a range from 300 to 800 microns is approximately 4.5 $\mu$m/hr. Due to fluctuations in the experimentally measured range expansion rate at different values of $a$ caused by errors in radius measurement, as well as the inherent challenges in capturing longer time lapses, it is challenging to discern such a minor variation in $\Delta a/\Delta t$. In particular, our uncertainty in $\Delta a/\Delta t$ is $\pm$2.5 µm when averaging over multiple values of $a$, which would put a change of 4.5 $\mu$m/hr within overlapping error bars (and, of course, our uncertainty will be higher if we do not average over multiple value of $a$.

### 4. Methods

| Strain | Species | Genotype | Gram+/- |
|---|---|---|---|
| C6706 | *V. cholerae* | Wildtype | - |
| C6706 | *V. cholerae* | $\Delta$ *vpsR* | - |
| C6706 | *V. cholerae* | $\Delta$ *hapR* | - |
| Z0R0001 | *A. veronii* | Wildtype | - |
| PA01 | *P. aeruginosa* | Wildtype | - |
| 0168 | *B. subtilus* | Wildtype | + |

**Table 1. Description of the bacterial strains used in this study. In the main text, we refer to *V. cholerae* $\Delta$ *vpsR* and *V. cholerae* $\Delta$ *hapR* as *V. cholerae (EPS-)* and *V. cholerae (EPS+)* respectively.**

#### A. Image analysis.

***A.1. Identification of biofilm center.*** After subtracting the background from the raw image using a best-fit polynomial, our next step in analyzing biofilm topographies involved precisely identifying the center of the bacterial colony. This step is particularly critical when calculating $L_e$ based on consecutive images. We initially identified points around the biofilm that were at approximately 5% of the maximum height and then determined the best-fitting circle using these points (Fig. S15). We tested three different threshold values: 5%, 10%, and 20% of the maximum height, which all yielded similar center positions with minimal differences in the coordinates and measured values. For example, the percent difference in volume, radius and height

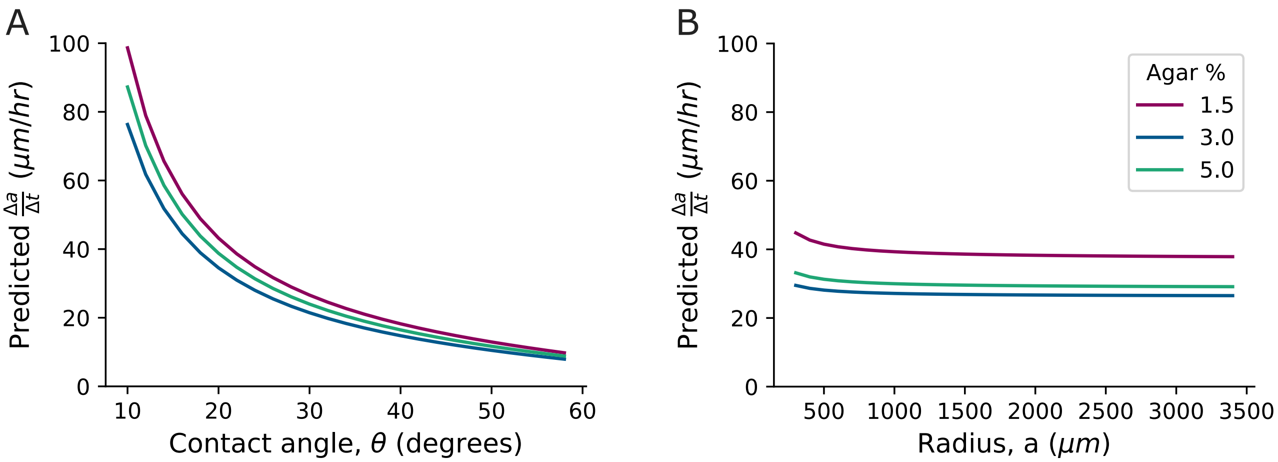


**Fig. S14.** **(A)** Predicted range expansion rate ($\Delta a/\Delta t$) is plotted against the contact angle for three different agar percentages. We used a fixed radius of 500 $\mu m$, the mean $L_e$ from three different replicates, and the mean doubling time $\tau$. Strong dependence of range expansion on contact angle is observed. **(B)** On the same scale, we plot the predicted range expansion rate against the colony radius. Consistent with the approximation made in Equation S35 and the predictions that follow, we observe that the range expansion rate saturates for larger values of the radius.

calculated from different thresholds is less than 0.5%. The center of this best-fit circle was used as the center of the biofilm. The center points calculated using this approach closely align with those determined using the coordinates based on the maximum height. However, for time lapse data sets, we used this approach rather than just picking the tallest point to be the center to safeguard against topographic fluctuations and measurement noise.

We tested three different threshold values: 5%, 10%, and 20% of the maximum height, all of which produced similar center positions with minimal differences in the coordinates. For example, the percent difference in volume, radius and height calculated from different thresholds is less than 0.5%.

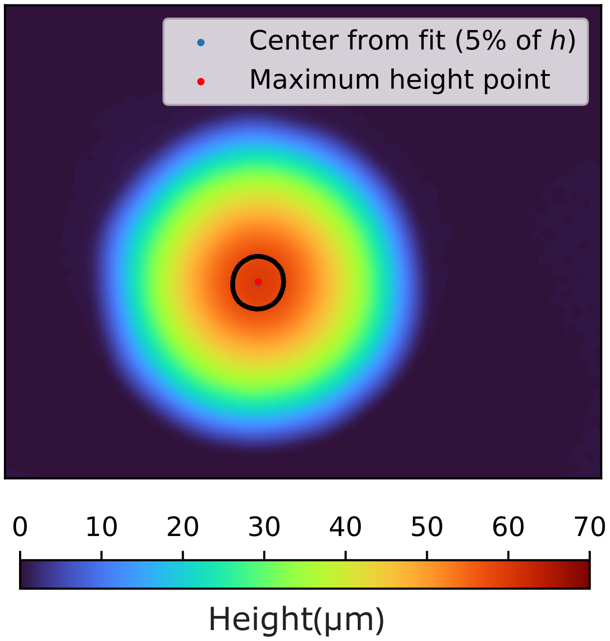


**Fig. S15.** A sample biofilm topography demonstrating the process of determining the center. We selected points around the center (denoted by the black circle) corresponding to 5% of the colony's maximum height and used these points to obtain a best-fit circle. The center obtained from the best-fit circle was used as the center of the colony. The center coordinates obtained through this process are robust against noise in the dataset and remain consistent across different threshold heights

***A.2. Contact angle measurement.*** We drew 20 radial lines from the center of the biofilm to the outer edge of the biofilm for each individual topographic image (See Fig. S16 A and B). Each line profile was then smoothed using the Savitzky-Golay filter to remove any data noise. We divided each line profile into approximately 40 different sections and determined the gradient along each of these sections. For each profile, the contact angle was calculated as the maximum gradient among all the sections in the line profile (See Fig. S16 C). Finally, the mean contact angle for the colony was computed by averaging the contact angles obtained from 20 different profiles around the colony.

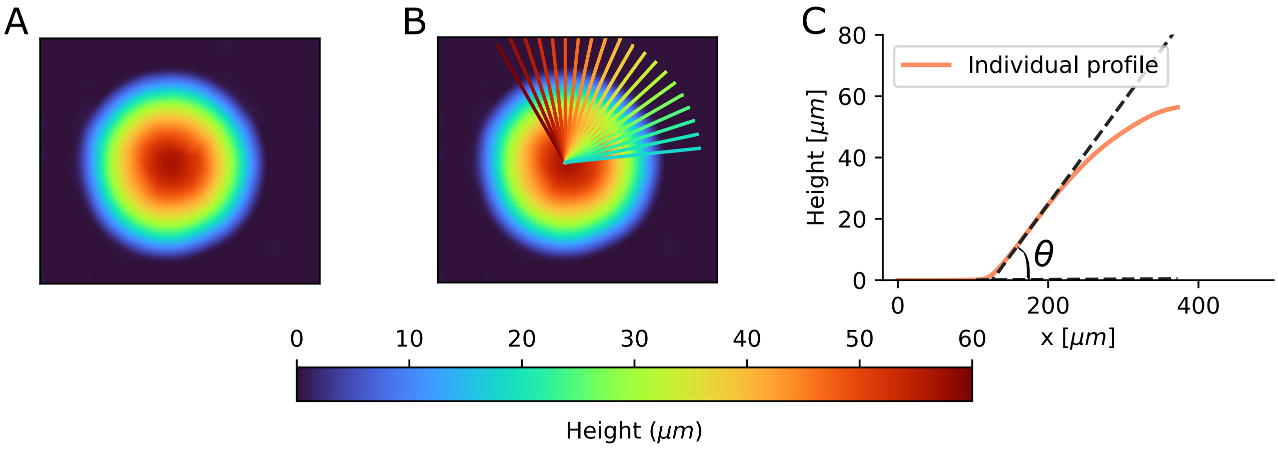


**Fig. S16. A)** A biofilm topography obtained through interferometry. **B)** To determine the contact angle, we obtained 20 line profiles from the center to the outer edge of the colony. **C)** For each line profile, we calculated the maximum gradient along the profile. The contact angle reported in the main text is the mean contact angle obtained from these 20 different line profiles.

1. EW Weisstein, Gaussian function (MathWorld–A Wolfram Web Resource) (2023).
